# Rescue of the Stargardt Disease phenotype in *Abca4* knockout mice through dietary modulation of the vitamin A receptor RBPR2

**DOI:** 10.1101/2025.06.27.662034

**Authors:** Rakesh Radhakrishnan, Matthias Leung, Drew Yochim, Heidi Roehrich, Scott W. McPherson, Glenn P. Lobo

**Affiliations:** Department of Ophthalmology and Visual Neurosciences, University of Minnesota, Lions Research Building, 2001 6th Street SE, Minneapolis, MN 55455, USA

**Keywords:** RBPR2, RBP4, Retinoic Acid, all-*trans*-retinol, Retinoids, HPLC, Retinoic Acid Response Elements, Promoter, β-carotene, lipofuscin, ABCA4, Stargardt Disease, RPE

## Abstract

Mutations in the *ABCA4* gene in Stargardt disease (STGD1) causes accumulation of cytotoxic lipofuscin, resulting in RPE atrophy and photoreceptor dysfunction. One component of lipofuscin is the all-*trans*-retinal derivative, bisretinoid *N-*retinylidene-*N-*retinylethanolamine (A2E). Since ocular A2E biosynthesis relies on circulating all-*trans*-retinol bound to retinol binding protein 4 (RBP4-ROL), we hypothesized that modulating vitamin A receptors, such as the retinol binding protein receptor 2, RBPR2, which regulate serum RBP4-ROL concentration, should attenuate A2E production. In-silico analysis revealed multiple retinoic acid response element (RARE) binding sites on the murine *Rbpr2* gene promotor, which was confirmed in vitro by EMSA and ChIP assays. In vitro luciferase assays showed that *Rbpr2* promotor activity was induced by exogenous β-carotene (BC) metabolites. Dietary BC supplementation of *Abca4^−/−^* mice, a mouse model for STGD1, increased hepatic all-*trans*-retinoic acid and 9-*cis*-retinoic acid production, which induced *Rbpr2* mRNA expression. This mechanism decreased serum RBP4 protein levels, fundus autofluorescence (AF) and ocular A2E accumulation, altogether improving photoreceptor and RPE function. Conversely, such a rescue was not observed in either *Abca4^−/−^* mice fed a diet devoid of BC or in double knockout *Rbpr2^−/−^*;*Abca4^−/−^* mice. Thus, there was a significant inverse correlation between dietary BC supplementation and *Rbpr2* gene presence in *Abca4^−/−^* mice, to that of lipofuscin accumulation in *Abca4^−/−^* mice on diets devoid of BC or in *Rbpr2^−/−^*;*Abca4^−/−^* mice. Our results provide impetus to pursue BC supplemented diets as therapeutic interventions for STGD1 patients with *ABCA4* gene mutations and identifies a novel role for the vitamin A receptor RBPR2 in this process.

## 1. INTRODUCTION

Stargardt disease (STGD1) is one of the most common forms of inherited retinal dystrophies, or inherited retinitis pigmentosa (RP) known, affecting approximately 1 in 10,000 people^1^. STGD1 is characterized as a cone-rod photoreceptor dystrophy, and differs from the typical presentation of RP in that the cones degrade prior to rod degeneration^2^. STGD1 is a childhood-onset disease, with a median age of onset at 8.5 years^3^ and typically presents initially as loss of bilateral visual acuity as a result of macular atrophy, a “darkened” choroid as a result of blockage of choroid fluorescence by accumulated lipofuscin, the presence of yellowish-white flecks of lipofuscin at the posterior pole of the retinal pigment epithelium (RPE) as seen through ophthalmoscopy or fundus autofluorescence (AF), disrupted visual function as measured through electroretinography, and RPE atrophy enlargement^3–8^.

The causative gene of STGD1 is the *ABCA4* gene, which encodes the <u>A</u>TP-<u>b</u>inding <u>c</u>assette subfamily <u>A</u> member <u>4</u> transporter (ABCA4)^7,8^. ABCA4 protein is expressed in the rim of the disc segments within photoreceptors, and functions as a flippase to transport accumulated *N*-retinylidene-phosphatidylethanolamine (*N*-ret-PE) outside the photoreceptor discs. *N*-ret-PE is metabolite resulting from the Schiff base formation between phosphatidylethanolamine (PE), a common phospholipid found within mammalian cell membranes, and the retinoid all-*trans*-retinal (atRAL), the consequential product after photoisomerization of the critical chromophore 11-*cis*-retinal in the visual cycle^9–11^. While most atRAL continues along the visual cycle through reduction to all-*trans*-retinol (ROL) through the action of retinol dehydrogenase 8 (RDH8), eventually leading to the regeneration of 11-*cis*-retinal chromophore, a large quantity of this atRAL reacts with PE to form the *N*-ret-PE metabolite^12–14^. Following the formation of *N*-ret-PE, another molecule of atRAL may react with *N*-ret-PE to form dihydropyridinium-A2PE, which subsequently undergoes a proton shift and proton elimination reaction to form A2PE. This can undergo a phosphate hydrolysis reaction catalyzed by phospholipase D to form *N*-retinyl-*N*-retinylidene ethanolamine (pyridinium bisretinoid-A2E). A2E is one of the main components of lipofuscin, a primary component of the previously mentioned yellowish-white flecks and contributes to the “darkened” choroid in STGD1. A2E is highly cytotoxic, and its accumulation in STGD1 has been directly attributed to its macular atrophic effects. There are several hypotheses as to how A2E exert its cytotoxic effects. First, bisretinoids such as A2E exhibit mild detergent like characteristics, which may disrupt membrane bilayers, including lysosomal membranes, potentially leading to disrupted proteolytic capabilities^15,16^. Second, the photodegradation products of A2E have been implicated in the degenerative effects associated with bisretinoid accumulation, including singlet oxygen species, epoxides, as well as oxo-aldehydes^14,15,17^.

A2E, being derived from atRAL originating from the visual cycle and its vitamin A/all-*trans*-retinol/ROL based intermediates, is therefore intrinsically linked to the broader system of dietary vitamin A homeostasis present in mammalian systems. Dietary obtained vitamin A, much like many other nutrients, must be maintained at homeostatic levels in an organism^10^. This is especially true for vitamin A/ROL since ROL is directly used to generate the critical light sensitive chromophore, 11-*cis*-retinal, that stimulates the photoreceptor to generate an electrical impulse. Consequently, modulating the biochemical components within serum ROL homeostasis could potentially influence the generation and accumulation of A2E containing lipofuscin within STGD1. Maintenance of serum ROL homeostasis requires an extensive system of receptors, binding proteins, specialized cells, and other biochemical mechanisms^10,11^. In the past, our laboratory has elucidated the role of the non-ocular vitamin A receptor, Retinol Binding Protein 4 Receptor 2 (RBPR2), in the maintenance of serum RBP4-ROL and whole-body ROL homeostasis within the murine system. RBPR2 is a relatively novel vitamin A receptor, and was identified in 2013 by the Graham laboratory^18^. RBPR2 is one of two known vitamin A receptors and functions to transport circulatory ROL into the cell when bound to its transporter Retinol Binding Protein 4 (RBP4-ROL)^18,19^. Through its expression within the liver, the main storage organ for retinoids, and its specific interactions with serum RBP4-ROL, RBPR2 likely serves as a key regulator for whole-body vitamin A homeostasis by modulating serum RBP4-ROL homeostasis^20^. Since lipofuscin/A2E generation is dependent upon the supply of ROL to ocular tissue via circulatory ROL bound RBP4 complex, which in turn is regulated in the liver by RBPR2, we have sought evidence that by modulating *Rbpr2* gene expression within the liver will affect the availability of RBP4-ROL precursors for ocular A2E production and accumulation in *Abca4^−/−^* mice, a mouse model of STGD1.

## 2. MATERIALS AND METHODS

### 2.1 Materials

All chemicals unless stated otherwise were purchased from Sigma-Aldrich (St. Louis, MO, USA) and were of molecular or cell culture grade quality. Cell culture reagents including Fetal Bovine Serum (FBS) were purchased from Gibco/ThermoFisher (Waltham, MA, USA). High Performance Liquid Chromatography (HPLC) standards were purchased from Toronto Research Chemicals (ON, Canada) or Sigma-Aldrich (St. Louis, MO, USA).

### 2.2 In-silico *Rbpr2* gene Promoter Analysis

We used NUBIScan version 2.0 software (http://www.nubiscan.unibas.ch/software), AliBaba 2.1 (http://www.gene-regulation.com), and EPD (Eukaryotic Promoter Database; https://epd.expasy.org/epd/) to identify putative nuclear receptor Retinoic Acid Receptor (RAR) and Retinoid X Receptor (RXR) binding sites on the mouse *Rbpr2* gene promoter region, as previously described by us^21^.

### 2.3 Dual-Luciferase Reporter Gene Assay

The murine *Rbpr2* promoter region (∼2.2 kb) was cloned using oligonucleotides listed in **Table 1** and inserted into the pGL3 luciferase reporter vector (Promega). 2 µg of purified plasmid DNA (*Rbpr2* promoter reporter vector or the empty pGL3 vector) was incubated with 4 µL of FuGENE HD Transfection reagent (Cat. No. E2311 Promega) for 15 min at RT and transfected individually in to COS1 cells (at ∼50% confluence) as per manufacturer’s protocol. At ∼12 h post-transfection, individually transfected COS1 cells were treated with either 2 µM all-*trans*-retinoic acid (RA), 2 µM 9-*cis*-retinoic acid, 2 µM β-carotene (BC), 2 µM 9-*cis*-retinol, 2 µM all-*trans*-retinol (ROL), or with vehicle control (DMSO). Post 24 h treatment the cells were harvested and the *Rbpr2* promoter activity in response to the individual retinoids was determined by Luciferase assay using the Pierce™ Renilla-Firefly Luciferase Dual Assay Kit (ThermoFisher). The cells were washed and harvested with 1X Lysis buffer and 20 mL loaded on a black opaque 96-well plate with 50 μL working solution; after incubating the mix at room temperature for 10-15 min, the luminescence was measured on a SpectraMax Gemini EM microplate spectrofluorometer (Molecular Devices). The data were normalized with blank and pGL3 basic vector samples and plotted in GraphPad Prism version 10.1.

**TABLE 1.**
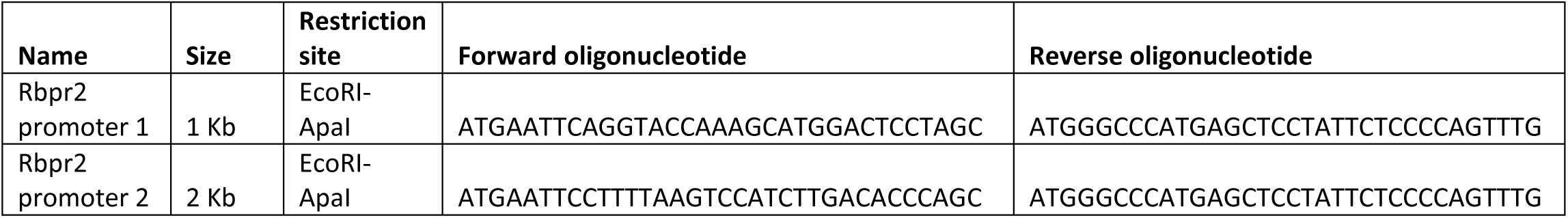
Oligonucleotide Sequences: Murine *Rbpr2* promoter cloning in the pGL3 plasmid.

### 2.4 Chromatin Immunoprecipitation Experiment

COS1 cells were cultured in DMEM medium with 10% FBS for five days. At ∼50% confluence, COS1 cells were transfected with 3 µg of pGL3-promoter plasmid containing the 2.2 kB *Rbpr2* promoter. Approximately 40 h post-transfection, cells were treated with either 2.5 µm ROL or 2.5 µm BC, for 24 h. Cells were then treated with 1% formaldehyde at 37^0^C with gentle swirling for 10 min to enable crosslinking of nuclear proteins with genomic DNA, followed by sonication to shear and obtain DNA fragment sizes between 150 and 250 bp in length, as described^22^. The Chromatin Immunoprecipitation (ChIP) assay was then performed as described by the manufacturer (Millipore), to evaluate binding of RARs and RXRs to the murine *Rbpr2* gene promotor. Approximately 4 µg of RARγ and RXRα antibody (Santa Cruz Biotechnologies or ThermoFisher) was used for immunoprecipitation separately, while 4 µg of IgG antibody was used as the negative control. Immunoprecipitated DNA was PCR amplified with the *Rbpr2* gene promoter oligonucleotides (ChIP-8 and ChIP-9) listed in **Table 2**. PCR products were electrophoresed on 2% agarose gels for visualization of amplified products.

**TABLE 2.**
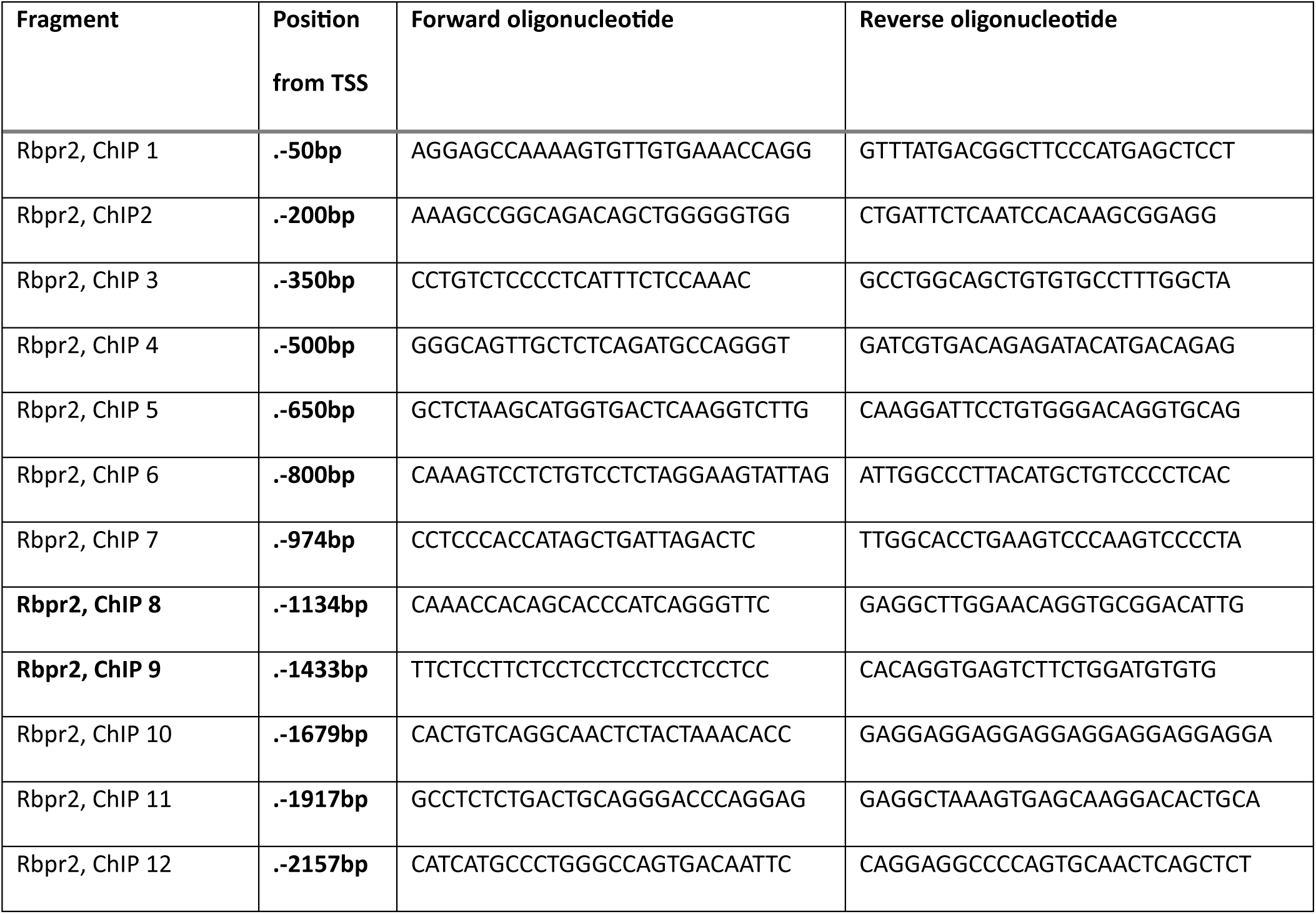
Oligonucleotide Sequences: Chromatin Immunoprecipitation (ChIP) Assay Only ChIP-8 and ChIP-9 oligonucleotide pairs were used for this assay.

### 2.5 Electrophoresis Mobility Shift Assay

The mouse *Rbpr2* promoter (∼2.2 kb) was subdivided into eight overlapping DNA fragments. Each fragment was PCR amplified from wild-type (WT) mouse genomic DNA prepared from liver with the oligonucleotides listed in **Table 3**. An aliquot of each PCR product was evaluated on 2% agarose gels to determine its size and purity. Electrophoresis Mobility Shift Assay (EMSA) was performed with SYBR™ Green & SYPRO™ Ruby EMSA stains (ThermoFisher). Liver extracts (∼5 µg) from C57BL/6 mice were immunoprecipitated with RXRα and RXRγ monoclonal antibodies and then incubated with individual *Rbpr2* promoter fragments (∼300 bp; 20 ng) for 1 h at 4°C. The protein-DNA complex was mixed with loading buffer and electrophoresed on 6% Tris-Borate-EDTA buffer PAGE gels. The DNA staining with SYBR Green and protein binding SYPRO Ruby staining signal images were captured sequentially as per manufacturer’s protocol using a ChemiDoc Imaging System (BioRad). The *Rbpr2* gene promoter fragments with various molecular weight complexes in the shifted bands were quantified (densitometry) using Image J software version 1.54. and normalized with only promoter DNA control band. All EMSA images in relevant figures are representative of three independent experiments.

**TABLE 3.**
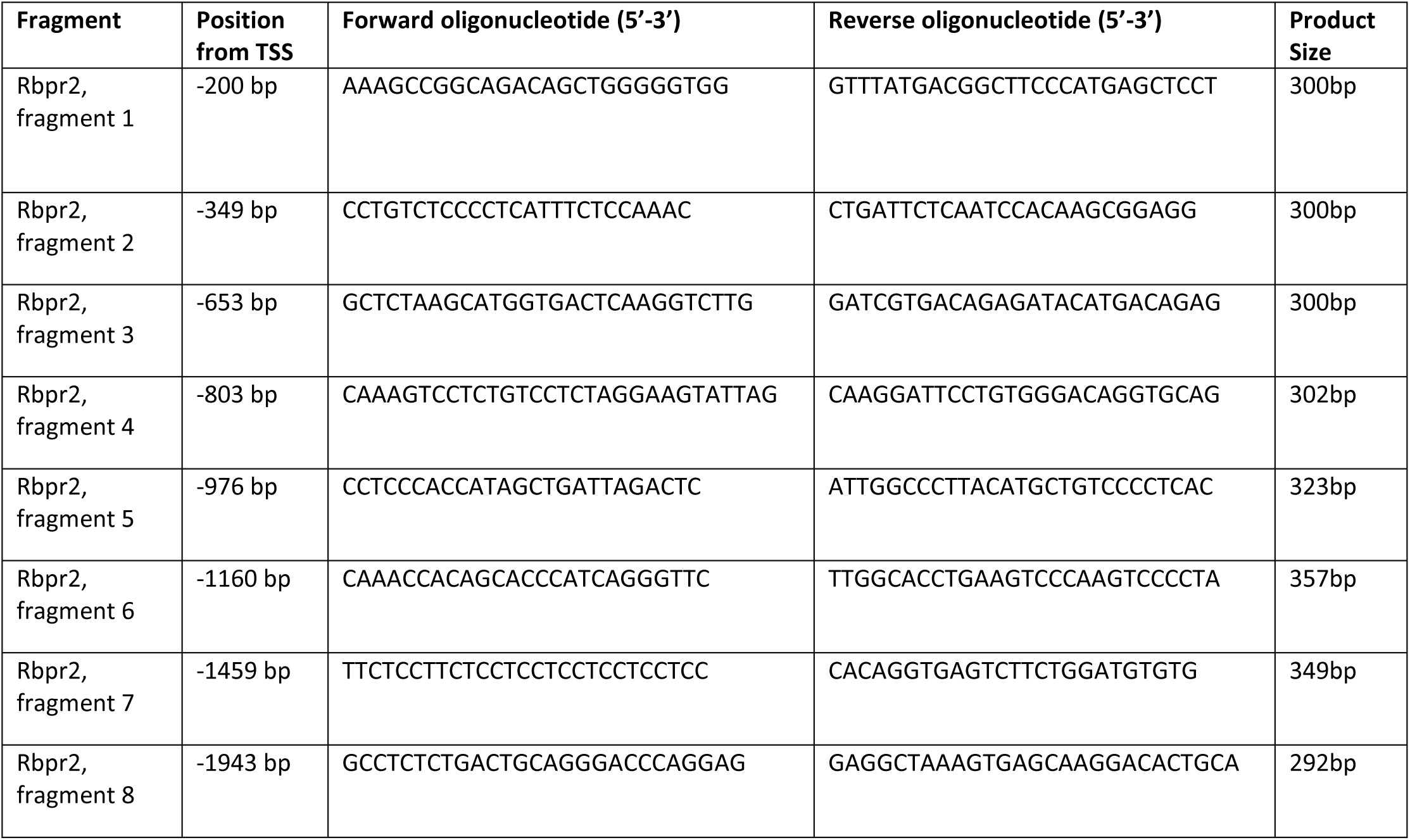
Electrophoresis Mobility Shift Assay (EMSA). Oligonucleotide sequences for generation of individual *Rbpr2* gene promotor fragments.

### 2.6 Animal handling and Carotenoid Feeding Experiment

Four-week-old WT mice (C57BL/6J) were purchased from Jackson (JAX) laboratories (RRID:IMSR_JAX:000664). *Rbpr2*-knockout (*Rbpr2^−/−^*) mice used in the study have been previously described^20^. *Abca4^−/−^* founders on a C57BL/6J background were purchased from Jackson laboratories (RRID:IMSR_JAX:026800). Breeding pairs and litters of *Rbpr2^−/−^*, *Abca4^−/−^*

, double knockout *Abca4^−/−^*;*Rbpr2^−/−^*, and WT mice were genotyped and found to be negative for the known *Rd8* and *Rd1* gene mutations, as previously described by us^20^. Breeding pairs were fed purified chow diets containing 8 IU of vitamin A/g (Research Diets, New Brunswick, NJ) and provided water ad libitum and maintained at 24°C in a 12:12 hour light-dark cycle. All animal experiments were approved by the Institutional Animal Care and Use Committee (IACUC) of the University of Minnesota (protocol #2312-41637A) and performed in compliance with the ARVO Statement for the use of Animals in Ophthalmic and Vision Research. Post weaning (P21), equal numbers of male and female mice were randomly distributed to chow diets containing 4 IU vitamin A/g (VAS diet) or β-carotene (BC supplemented diet) feeding groups. For experiments, WT, *Abca4^−/−^*, or double knockout *Abca4^−/−^*;*Rbpr2^−/−^* mice (both males and females) were fed purified rodent diets (AIN-93G; Research Diets, New Brunswick, NJ) containing the recommended 4 IU of vitamin A/g (VAS) or β-carotene (BC) supplemented diets containing 0.150 mg/g based on the AIN-93G diet (Research Diets, New Brunswick, NJ) for up to 6-months.

### 2.7 Optical Coherence Tomography and Fundus Imaging

Mice from various cohorts were sedated with 5% isoflurane with adjustable room airflow from Kent Scientific’s mouse anesthesia machine. After sedation, the isoflurane was reduced to 2%, and the mouse eyes were dilated with a 1:1 mix of Tropicamide 1.0% Ophthalmic Solution (Cat. No. NDC 61314-355-02 Sandoz) and Phenylephrine HCl 2.5% Solution (Cat. No. NDC 17478-201-15 Akorn). Alcon Systane Lubricant Eye Gel was applied to moisturize the eye, and the Phoenix Micron IV image guided OCT2 instrument, equipped with a specialized objective lens, was used to capture OCT images. After capturing the OCT images, the lens of the instrument was replaced with a fundus-contact lens, and fundus images and autofluorescence of the fundus (AF) was captured. The A2E lipofuscin in-vivo excitation spectra is in range of 400-590nm^23^, the instrument excitator filter was opened for the excitation wavelength range on the fundus and barrier filter on the instrument for fluorescence was used to capture emission wavelength.

### 2.8 RNA Isolation and Quantitative Real-Time Reverse Transcription PCR

After euthanizing mice, total RNA from ∼200 mg liver samples were extracted using a total RNA isolation kit (Qiagen). Four µg of isolated RNA with purity 260/280 nm ratio of 2.0 was reverse transcribed with an RNA to cDNA system (Applied Biosystems). An aliquot of cDNA (∼1 µL from a 20 µL reaction mix) was then subjected to qRT-PCR using TaqMan™ Fast Advanced Master Mix and ThermoTaq probes for *Rbpr2* (GenBank: AK004855.1) and *Gapdh* genes (GenBank: AK002273.1). The reaction was performed in a BioRad qPCR instrument and quantified using the ΔΔCt method. The ΔCt value was calculated by (Ct value (gene of interest) – Ct value (housekeeping gene)), then ΔΔCt was calculated by (ΔCt value (BC diet)-ΔCt value (VAS diet)). Then the fold change was calculated by 2-ΔΔCt. All qRT-PCR experiments were performed in triplicate and repeated thrice, using freshly synthesized cDNA. *Rbpr2* qRT-PCR values were normalized to 18S RNA, and the ΔΔCt method was employed to calculate fold changes. Data of qRT-PCR were expressed as mean ± standard error of mean (SEM).

### 2.9 High-Performance Liquid Chromatography analyses of retinoids

Retinoid isolation procedures were performed under a dim red safety light (600 nm) in a dark room, as previously established by us^20^. Animals were first euthanized with CO_2_ asphyxiation, and pertinent tissues were removed from the carcass. Various murine tissues (liver, serum, kidney) were homogenized separately in 0.9% saline with a handheld tissue grinder, consisting of a glass tube and glass pestle. Methanol (2 mL) was added to the homogenate to precipitate the proteins. The retinoid content from the homogenate was then extracted with 10 mL of hexane (twice), with the aqueous layer subsequently removed. The combined hexane extracts were then evaporated with a vacuum evaporator, re-suspended in 100 μL of hexane, then injected into an HPLC for analysis. HPLC analysis was performed on an Agilent 1260 Infinity HPLC with a UV detector. The HPLC conditions employed two normal-phase Zorbax Sil (5 μm, 4.6 × 150 mm) columns (Agilent, Santa Clara, CA, USA), connected in series within the Multicolumn Thermostat compartment. Chromatographic separation was achieved by isocratic flow of mobile phase containing 1.4% 1-Octanol/2% 1,4-Dioxane/11.2% Ethyl Acetate/85.4% Hexane, at a flow rate of 1 ml/min for 40 min. Retinaldehydes, retinol, and retinyl esters were detected at 325 nm using a UV-Vis DAD detector, while the UV absorbance spectra was collected from 200 nm – 700 nm. For quantifying molar amounts of retinoids, the HPLC was previously calibrated with synthesized standards and as previously described by us^20,23^. Calculation of concentration (µM): Standards were injected in concentrations ranging from 0-3.5 µM prepared solutions in the mobile phase. The plotted concentrations were fit through linear regression to obtain R-equation (y=mx+c) where *y* is the peak area (mAU*sec), *m* is the slope of the calibration curve, and *c* is the y-intercept. The area from the HPLC peaks of the samples were interpolated into concentration and expressed as picomoles. For eyes the values are expressed as picomoles/eye; for liver and kidney the values are expressed as picomoles/mg of tissue; for serum the ROL values are expressed as picomoles/ microliter.

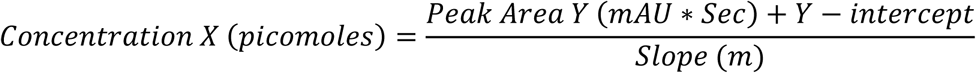

### 2.10 Biomimetic Synthesis of Bisretinoid Standards

Bisretinoids were synthesized by incubating 32 µg ethanolamine and approximately higher equivalent of ∼0.3 mg or more quantity of all-*trans*-retinal in 3 mL of ethanol stirred and incubated at room temperature for 3-7 days in the dark^24^.

### 2.11 Spectrophotometric quantification of synthesized bisretinoid standards and Standard Curve Generation

To quantify the concentration of synthesized bisretinoid standard, we first separated and resolved the bisretinoids utilizing the HPLC method described below and collected the bisretinoid fractions using an HPLC fraction collector. Utilizing the Beer-Lambert Law and the molar absorptivity of the pertinent bisretinoid, we quantified the concentration of the collected fractions. The molar absorptivity coefficient for A2E in methanol at 439 nm is 36,900, while the molar absorptivity coefficient for isoA2E in methanol at 426 nm is 31,000^24^.

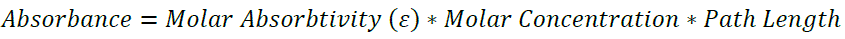

Standards were then injected in amounts ranging from 250-2500 picograms prepared solutions in the mobile phase. The plotted concentrations were fit through linear regression to obtain R-equation (y=mx+c) where *y* is the peak area (mAU*sec), *m* is the slope of the calibration curve, and *c* is the y-intercept. The area from the HPLC peaks of the samples (mAU*sec) are interpolated into concentration and expressed as picomoles. For eyes, the values are expressed as picomoles/ eye.

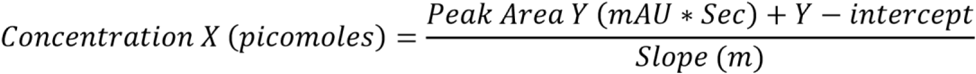

### 2.12 Bisretinoid extraction from Murine Eyes

All bisretinoid extractions were performed in a dark room illuminated by dim red light (600 nm) to minimize potential photoisomerization. Mice were euthanized through CO_2_ asphyxiation, and whole eye globes were removed and homogenized in 0.5 mL Chloroform / 0.5 mL Methanol / 0.25 mL 1X PBS with a manual tissue grinder. The homogenate was then centrifuged to separate out the supernatant. The supernatant was removed and vacuum evaporated and resuspended in 0.1% TFA in methanol. This mixture was centrifuged again, and the final supernatant injected in to the HPLC for analysis.

### 2.13 High Performance Liquid Chromatography analysis and Quantification of Bisretinoids in Murine Eyes

High Performance Liquid Chromatography (HPLC) analysis was performed on an Agilent 1260 Infinity HPLC with a UV/Vis detector. The ZORBAX Eclipse Plus C18 (4.6 x 250 mm, 5 µm) column (Agilent, Santa Clara, CA, USA) was used to resolve bisretinoids, with the column compartment set to 20°C. Chromatographic separation was achieved using a gradient flow facilitated by a binary pump system, with acetonitrile in one pump, and 0.1% TFA in water in the other. The gradient flow was as follows: 50% A and 50% B (0-4 min), 20% A and 80% B (4-6 min), 10% A and 90% B (6-10min), 100% B (10-20 min). The flow rate was set to 0.5 mL/min for the duration of the run. Bisretinoids were best detected at 440 nm using a UV-Vis DAD detector, while the UV absorbance spectra was collected from 200 nm – 700nm.

### 2.14 Confocal Microscopy for Lipofuscin/ A2E granule visualization in murine RPE

Eye globes from mice in the respective cohorts were harvested and fixed in Davidson’s fixative overnight. The globes were then embedded in OCT compound (Tissue-Tek) and flash frozen. 10 microns thick tissue slices were sectioned with a microtome and layered on to a microscope glass slide. The slides were washed with 1X PBS then mounted with VectaShield Antifade Mounting Medium with DAPI for nuclear staining. Images were captured using 60X water immersion objectives. The A2E autofluorescence emission spectrum, spanning a continuous wavelength range from 520 to 800 nm, was detected using Laser settings of 445 nm and 405 nm for A2E excitation, resulting in emission filter signals at 570-616 nm and 607-850 nm. Additionally, 405 nm was used for the DAPI nuclear staining emission signal, with a wavelength of ∼460 nm^23,26^. The images were processed and calibrated with the ZEISS ZEN 3.4 software package. The autofluorescence signal in RPE layer from all the groups were quantified using Image J or Fiji (NIH) version 1.54, and the data-plotted in GraphPad Prism version 10.3.1.

### 2.15 Western Blot Analysis and Densitometry for Protein Quantification

Total protein from mouse serum was solubilized using RIPA lysis buffer containing protease inhibitors (Roche, Indianapolis, IN). Approximately 25 µg of total protein was electrophoresed on 15% SDS-PAGE gels and transferred to PVDF membranes. Membranes were stained with Ponceau S and images captured. The blots were then washed, destained, and probed with primary antibodies against RBP4 (1:250, Polyclonal Rabbit anti-human RBP4) in buffer (0.2% Triton X-100, 2% BSA, 1X PBS). HRP conjugated secondary antibodies (BioRad) were used at 1:5,000 dilution. Protein expression was detected, and images captured using a ChemiDoc Imaging System (BioRad). The relative intensities of each protein band were quantified (densitometry) using Image J software version 1.54 and normalized to loading control (Ponceau S staining) images. Each western blot analysis was repeated thrice.

### 2.16 Immunohistochemistry and Fluorescence Imaging

Mice, at 7-months of age in the different cohorts, were euthanized by CO_2_ asphyxiation and cervical dislocation. Eyes were enucleated and fixed in Davidson’s fixative at RT for 24 h. Paraffin-embedded retinal sections (∼10 microns) were processed for antigen retrieval and immunofluorescence. Primary antibodies: anti-rhodopsin 1D4 for mouse rod opsin (1:200; ab5417 Abcam), anti-Red/Green Opsin Polyclonal Antibody for mouse cone opsin (1:200; OSR00222W ThermoFisher Scientific) were diluted in blocking solution for overnight incubation with the samples. All secondary antibodies (Alexa Fluor 488 or Alexa Fluor 594) were used at 1:5000 concentration (Molecular Probes). 4′,6-diamidino-2-phenylendole DAPI; VECTASHIELD® Mounting Medium (H-1200 Vector Laboratories) was used to label nuclei. The retinal section Z-stack images were obtained using a 60X objective and sectioning on the Keyence BZ-X800 microscope. The images were processed using the Z-projection Max Intensity method. The fluorescence intensities and various length measurements of retinal sections were quantified using ImageJ or Fiji (NIH), and the data were plotted in GraphPad Prism.

### 2.16 Statistical Analysis

Data were expressed as means ± standard error mean, statistical analysis by ANOVA and Student t-test. Differences between means were assessed by Tukey’s honestly significant difference (HSD) test. P-values below 0.05 (P<0.05) were considered statistically significant. Statistical analysis was carried out using GraphPad Prism v 10.1.

## 3. RESULTS

### 3.1 In-silico analysis of murine *Rbpr2* promotor region reveals putative RARE binding sites

RARs and RXRs function by binding to specific retinoic acid response elements (RAREs) within regulatory/ promoter regions of target genes. We used NubiScan Version 2.0 and AliBaba 2.1 software to identify putative RAREs in the murine *Rbpr2* gene promoter, as achieved previously by us for other gene promoters^21,22^. These programs predicted multiple direct repeat (DR-2) RAREs sites located upstream of the ATG start codon of the *Rbpr2* gene promoter region (**Supplementary Figure S1A**). Additionally, promoter analysis from the eukaryotic promoter database UCSC genome browser for the presence of RAR and RXR binding elements further confirmed presence of RARE regulatory elements on the murine *Rbpr2* promoter (**Supplementary Figures S1B and S1C**).

### 3.2 RAR and RXR interacts with the murine *Rbpr2* promoter region in Chromatin Immunoprecipitation Assays

To investigate whether RARs and RXRs associate with the DR-2 RARE sites on the murine *Rbpr2* gene promoter region, we performed chromatin immunoprecipitation (ChIP) assays in COS1 cells using antibodies specific for RARγ and RXRα and *Rbpr2* gene oligonucleotide pairs designed for detection of the putative RARE sites in immune-precipitated DNA fractions (**Figure 1A**). Here we observed that PCR products were clearly detectable in chromatin fractions immunoprecipitated with individual RARγ and RXRα antibodies (**Figure 1B**). In contrast, no PCR products were detected when IgG antibody was used as a control in this assay (**Figure 1B**). Thus, our analysis provides evidence that RARs and RXRs bind to the computer predicted RARE elements within the murine *Rbpr2* gene promotor.

**Figure 1:**
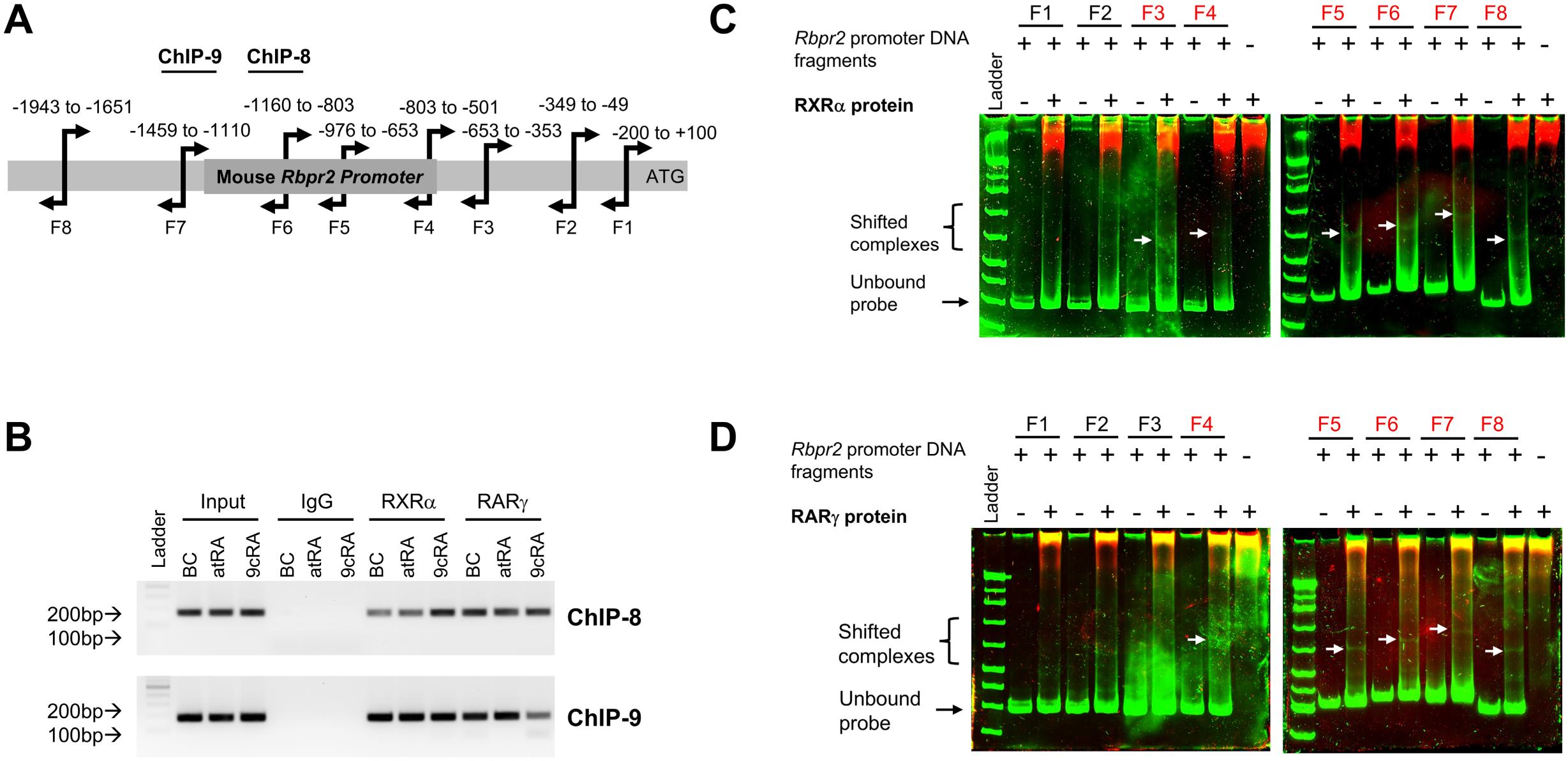
ChIP and EMSA assays identify multiple RAR and RXR binding sites on the murine *Rbpr2* gene promoter. (**A**) Schematic representation of ∼2.2 kb of the mouse *Rbpr2* promoter. (**A**) For electromobility shift assays (EMSA), genomic DNA was isolated from a wild-type/WT mouse liver and using oligonucleotide pairs listed in Table 3, the full-length *Rbpr2* gene promoter was PCR amplified into eight overlapping fragments (F1-F8) as indicated. Individual fragment sizes and positions are relevant to the *Rbpr2* gene ATG start site. Position of the two separate oligonucleotide pairs used in the chromatin immunoprecipitation (ChIP) assay are indicated (ChIP-8 and ChIP-9). (**B**) To show binding of RAR and RXR to the RARE sites on the mouse *Rbpr2* promoter, COS1 cells were transfected with pGL3 vector containing the full-length *Rbpr2*-promoter fragment (∼2.2 kb). After 40 h transfection, COS1 cells were incubated with either 2 µM all-*trans*-retinoic acid (atRA), β-carotene (BC), or 9-*cis*-retinoic acid (9cRA) for 24 h. ChIP assays in COS1 cells were performed with commercial anti-RARγ and anti-RXRα antibodies. The 10% of initial cell extract was used as input and IgG antibody was used as a control in the pull-down. Two oligonucleotide pairs flanking the F6 and F7 *Rbpr2* gene promoter region was designed for its detection in precipitated DNA fractions, namely ChIP-8, oligonucleotide pair 1, and ChIP-9, oligonucleotide pair 2, as shown in panel A. In EMSAs, individual *Rbpr2* gene promoter fragments were incubated with either RXRα (**C**) or RARγ (**D**) protein. *Rbpr2* promoter fragments F3-F8, each capable of interacting with RXRα or RARγ protein is indicated by the complex/shift (white arrows) and faster migrating unbound DNA probes are shown as free probes by the black arrow. Protein-DNA complexes were resolved on 6% PAGE gels, and results were obtained by Chemiluminescence. Shifts of the RAR and RXR protein bound *Rbpr2*-probe complexes are indicated by arrows. Although the free unbound probe migrated rapidly upon electrophoreses, the shifted RAR-*Rbpr2* and RXR-*Rbpr2* bound probe migrated slower and above the labeled free DNA probe. The figures show representative images from at least three independent experiments.

### 3.3 Electrophoresis Mobility Shift Assays of the *Rbpr2* promoter reveals multiple RARE binding sites

To confirm the observations obtained from the In-silico computer analysis and ChIP assay above, we used Electrophoresis Mobility Shift Assays (EMSA) to determine if a physical interaction exists between RAR and RXR and the RARE sites on the murine *Rbpr2* promotor. EMSA assays with individual *Rbpr2* promotor DNA fragments (**Figure 1A**) and RARγ and RXRα protein revealed that both RAR and RXR bound six of the eight *Rbpr2* gene promotor fragments in COS1 cells, wherein slower DNA-protein migration patterns were observed (arrows in **Figures 1C and 1D**). Thus, our EMSA analysis provides further evidence that RAR and RXR can physically bind to the distal RARE sites on the promoter region of murine *Rbpr2* gene.

### 3.4 Luciferase assays of mouse *Rbpr2* promoter reveals induction by exogenous carotenoids

We next tested the in vitro effects of various retinoids and carotenoids on murine *Rbpr2* gene promoter activity using luciferase assays. To achieve this, a full-length ∼2.2 kb region of the murine *Rbpr2* promoter region was PCR amplified using gene specific oligonucleotides and cloned into the pGL3 luciferase reporter plasmid, as outlined in the methods section. The pGL3-*Rbpr2* plasmid or empty pGL3 plasmid was transfected into COS1 cells and then treated individually with retinoids, all-*trans*-retinoic acid (atRA), 9-*cis*-retinoic acid (9cRA), 9-*cis*-retinol (9cROL), all-*trans*-retinol (atROL), and carotenoids (β-carotene). The influence of individual retinoids and carotenoids on *Rbpr2* gene promoter activity was determined by Luciferase assay. As expected, the empty pGL3 plasmid transfected COS1 cells, empty pGL3 transfected COS1 cells with atRA, and empty pGL3 transfected COS cells with vehicle (DMSO), showed no luciferase activity (**Supplementary Figures S2A and S2B**). Interestingly, we observed that pGL3-*Rbpr2* expressing cells treated with exogenous BC showed a significant induction of *Rbpr2* mRNA activity, while the individual retinoids (atRA, 9cRA, atROL, and 9cROL) treated cells caused a suppression of *Rbpr2* promoter activity (**Supplementary Figures S2A and S2B**). To determine the mechanism for these observations we isolated total retinoids from individual COS1 treated cells and subjected them to HPLC analysis to determine the retinoid profiles. As expected, no retinoids were detectable in control COS1 cells (empty pGL3 plasmid and treated with DMSO) (**Supplementary Figure S2C**). Conversely, cells expressing pGL3-*Rbpr2* gene plasmid and treated with atROL or BC showed a retinoid profile on HPLC analysis. Herein, we observed that exogenous BC treated cells showed significantly higher production and concentration of retinoic acid isomers (atRA, 9cRA, and 13cRA), compared to the all-*trans*-ROL treated cells (**Supplementary Figure S2D vs. S2E;** individual retinoic acid isomers quantified and shown in **Supplementary Figures S2F-S2H**).

### 3.5 Dietary β-carotene induces hepatic *Rbpr2* expression and modulates serum RBP4-ROL levels in wild-type mice

Based on our in vitro observations above, we tested whether a similar mechanism could exist in vivo. For this, cohorts of 4-week-old wild-type (WT) mice were fed a diet supplemented with BC or a diet with vitamin A/ retinyl palmitate (VAS), the latter a source for all-*trans*-retinol. After 3-months of feeding, mice were euthanized, and their liver and serum were collected. Quantitative PCR (qRT-PCR) analysis showed that hepatic *Rbpr2* mRNA levels were significantly induced (∼5.4 fold) in WT mice fed a BC supplemented diet, compared to those fed a VAS diet (**Supplementary Figure S3A**). Western blot analysis showed that serum RBP4 protein levels were significantly lower in WT mice fed a BC diet, compared to those fed a VAS or chow diet (**Supplementary Figure S3B** and quantified in **S3C**). HPLC analysis for hepatic retinoids showed a similar trend to those observed above in COS1 cells, wherein, we observed an increased production and higher concentration of retinoic acid isomers (atRA, 9cRA, and 13cRA) in the liver of BC supplemented WT mice, compared to those fed a VAS diet (**Supplementary Figure S3D vs. S3E,** and quantified in **Supplementary Figures S3F-S3H**). Taken together, our in vitro and in vivo results support the observation that exogenous BC in COS1 cells and dietary BC supplementation in WT mice can induce *Rbpr2* mRNA expression and activity via increased RA synthesis.

### 3.6 β-carotene supplementation of *Abca4^−/−^* mice increases hepatic retinoic acid production and induces *Rbpr2* mRNA expression

Since all-*trans*-RAL is a primary reactant in the biogenesis of ocular lipofuscin/A2E, we tested the strategy identified above of reducing serum RBP4 availability in *Abca4^−/−^* mice^25–28^. This we hypothesized would limit serum ROL precursors for ocular all-*trans*-RAL formation in mouse models that can accumulate cytotoxic lipofuscin/A2E. To achieve this, we obtained *Abca4^−/−^*mice and crossed them with *Rbpr2^−/−^* mice. Then, individual cohorts of age-matched WT, *Rbpr2^−/−^*, *Abca4^−/−^*, and the double knockout *Rbpr2^−/−^*;*Abca4^−/−^* mice were fed a vitamin A sufficient/VAS diet (without BC) or a BC supplemented diet, starting at 1-month to 7-months of age. At 1-month of age, we collected blood to quantify serum RBP4 protein. After 3-months (4-month-old mice) and at 6-month (7-month-old mice) of feeding, mice in the individual cohorts were subjected to non-invasive tests for visual function and then euthanized, and their tissues and serum were collected. As we observed above in WT animals, qRT-PCR analysis showed that hepatic *Rbpr2* mRNA levels were significantly induced and upregulated (∼6.2 fold) in BC supplemented *Abca4^−/−^*mice, compared to VAS *Abca4^−/−^* mice (**Figure 2A**). Western blot analysis showed that serum RBP4 protein levels were significantly lower in BC supplemented *Abca4^−/−^* mice, compared to VAS *Abca4^−/−^*mice, at the 7-month of age (**Figures 2B and 2C**). Interestingly, as these mice age, serum RBP4 protein levels continued to rise in VAS *Abca4^−/−^*mice, conversely, it was observed to be lower in BC supplemented *Abca4^−/−^* mice (**Figures 2B and 2C**; 1-month-old vs. 7-month-old mice). HPLC analysis for hepatic retinoids in BC supplemented *Abca4^−/−^* mice showed a similar trend to those observed in WT mice. Together this data showed increased production and higher concentration of BC and retinoic acid isomers (atRA, 9-*cis*-RA, and 13-*cis*-RA) in the livers of BC supplemented *Abca4^−/−^* mice, compared to VAS *Abca4^−/−^*, *Rbpr2^−/−^*, or double knockout *Rbpr2^−/−^*;*Abca4^−/−^*mice (**Figure 2D vs. 2E,** individual RA isomers quantified in **Figures 2F-2K**, and BC quantified in **Supplementary Figure S4**).

**Figure 2:**
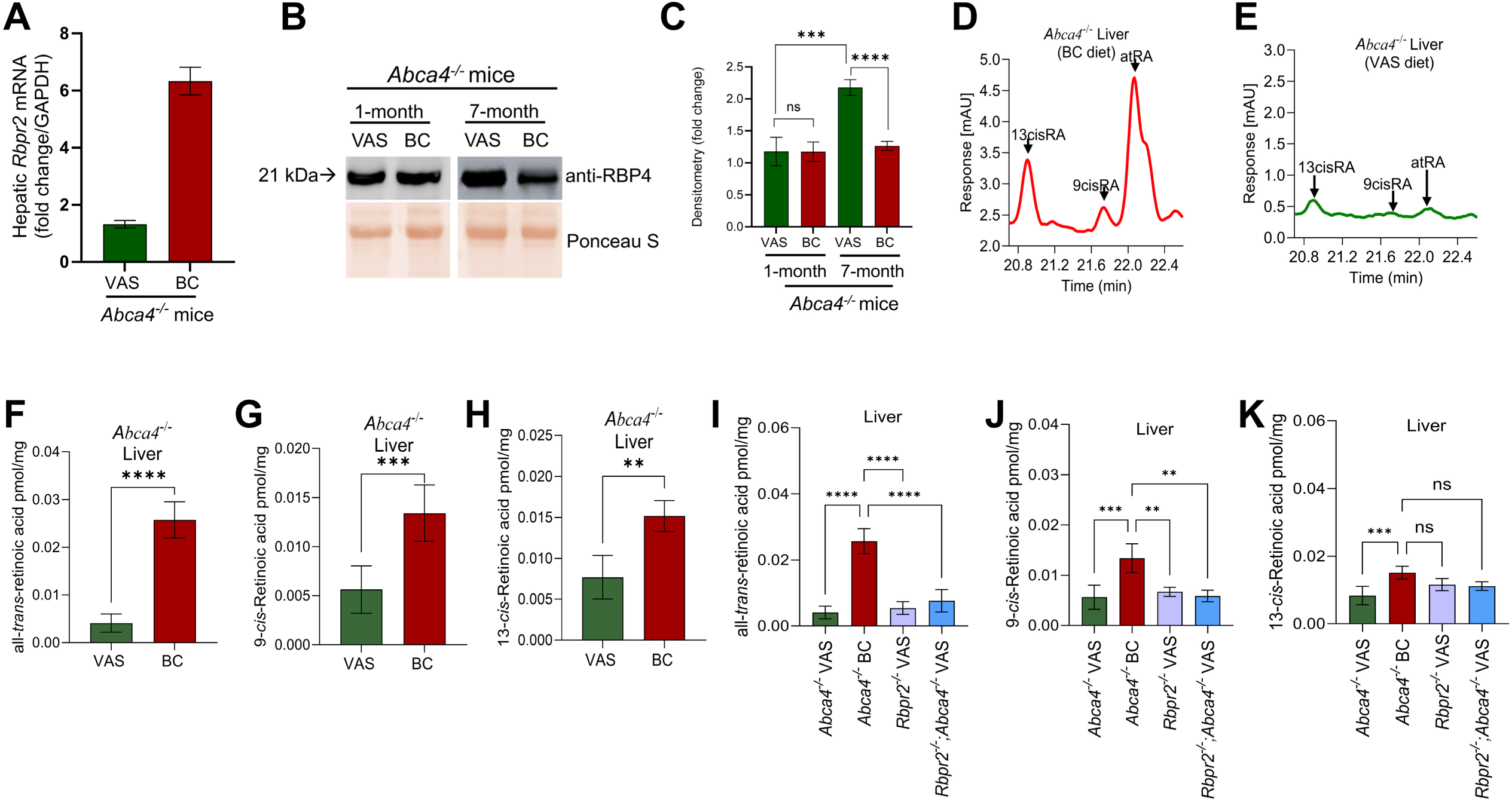
Long-term dietary β-carotene supplementation of *Abca4^−/−^*mice induces hepatic *Rbpr2* mRNA expression and increases hepatic retinoid production. (**A**) Quantitative RT-PCR analysis of *Rbpr2* mRNA expression in liver of *Abca4^−/−^*mice at 7-months of age, fed either a VAS or BC supplemented diet. (**B**) RBP4 protein expression in serum of *Abca4^−/−^* mice fed a VAS diet or BC supplemented diet, determined by western blot analysis and (**C**) protein quantified by densitometry. Ponceau S was used to confirm equal protein loading among samples. Approximately 50 µg of protein was loaded. The 1-month timepoint reflects start of the dietary intervention (either VAS or BC). (**D**, **E**) HPLC analysis of hepatic retinoic acid isomers extracted from *Abca4^−/−^* mice fed either a BC supplemented diet or a diet devoid of BC (VAS diet), respectively. (**F-K**) Quantification of various retinoic acid isomers in liver of mice from different cohorts fed with dietary BC or a diet devoid of BC (VAS diet). VAS, vitamin A sufficient diet; BC, β-carotene supplemented diet. n=4-6 mice per cohort and dietary condition. atRA, all-*trans*-retinoic acid; 9cisRA, 9-*cis*-retinoic acid; 13cRA, 13-*cis*-retinoic acid. Student t-test; *P<0.05; **P<0.01. n=4-6 mice per cohort.

### 3.7 Longitudinal recordings of 488 nm Fundus Autofluorescence intensity in *Abca4^−/−^* mice

A feature of *ABCA4*-mediated retinal and macular dystrophies is the accumulation of auto fluorescence (AF) lipofuscin/A2E pigments in the RPE^29^. We therefore investigated the changes in fundus AF levels over time in WT, *Abca4^−/−^*, *Rbpr2^−/−^*, and *Rbpr2^−/−^;Abca4^−/−^* mice, previously fed a BC supplemented or a diet devoid of BC (VAS diet). Repeated fundus recordings in the same animal cohorts were performed at 4-months of age (3-months post dietary BC supplementation or VAS diet) and then at 7-months of age (6-months post dietary BC supplementation or VAS diet). In control VAS WT and BC supplemented WT mice, 488 nm fundus AF was not detected at either 4– or 7-months of age (**Figures 3A and 3B**), likewise no AF was detected in adult VAS *Rbpr2^−/−^* mice at 7-months of age, indicating that loss of the vitamin A receptor *Rbpr2^−/−^* does not result in fundus AF lipofuscin accumulation (**Figure 3B**). At 4-months of age, fundus AF in double knockout *Rbpr2^−/−^;Abca4^−/−^*mice was significantly higher to that in VAS *Abca4^−/−^* mice, indicating that genetic loss of *Rbpr2* can accelerate AF accumulation in the fundus of these mice (**Figures 3A and 3A’**). Thereafter, 488 nm fundus AF rose significantly in both *Abca4^−/−^*and double knockout *Rbpr2^−/−^;Abca4^−/−^* mice fed a VAS diet, compared to either WT or BC supplemented *Abca4^−/−^* mice (**Figure 3A vs 3B**).

**Figure 3:**
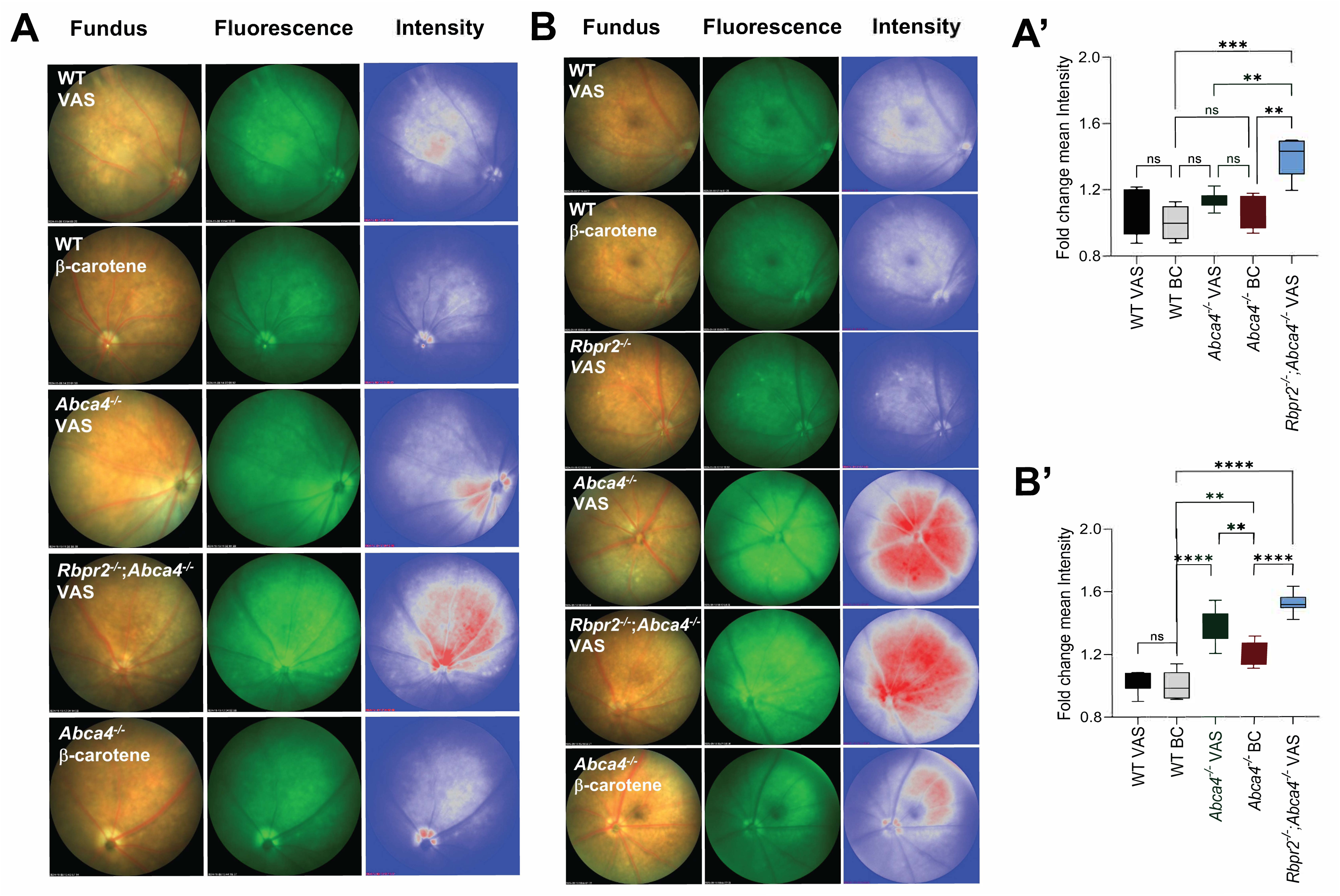
Long-term dietary β-carotene supplementation of *Abca4^−/−^*mice reduces fundus autofluorescence (AF) intensity. Repeated fundus recordings in the same animal cohorts were performed at (**A**) 4-months of age (3-months post dietary supplementation) and then at (**B**) 7-months of age (6-months post dietary supplementation). Shown are the changes in fundus autofluorescence (AF) levels over time in WT, *Abca4^−/−^*, *Rbpr2^−/−^*, and double knockout *Rbpr2^−/−^;Abca4^−/−^* mice fed either a β-carotene supplemented diet or a diet devoid of any β-carotene (VAS diet), which were evaluated at 488 nm. The relative intensity comparison was performed via pseudo-color Look Up Table on 8-bit images in Image J, showing the distribution of autofluorescence intensities in Fundus images. Red represents high intensity and Blue represents low intensity AF, in the respective 4– and 7-month cohorts. (**A’**) Fold change mean intensity of AF levels among cohorts at 4-months of age, and (**B’**) fold change mean intensity of AF levels among cohorts at 7-months of age. n=4-6 mice per cohort. VAS, vitamin A sufficient diet; BC, β-carotene supplemented diet. ANOVA, **P<0.01; ***P<0.005; ****P<0.001.

### 3.8 Comparison of Fundus Autofluorescence recordings in *Abca4^−/−^* mice

A2E is regarded as a major fluorophore of lipofuscin in the ocular fundus^25–28^. We therefore assessed if A2E levels in *Abca4^−/−^* and *Rbpr2^−/−^;Abca4^−/−^*mice fed a VAS diet increased in parallel with 488 nm autofluorescence (AF) levels, in comparison to BC supplemented *Abca4^−/−^* mice. AF ratios between *Abca4^−/−^* and *Rbpr2^−/−^;Abca4^−/−^*fed a VAS diet and BC supplemented *Abca4^−/−^* mice confirmed results from the longitudinal data set. At the 4-months of age timepoint, 488 nm AF levels in the double knockout *Rbpr2^−/−^;Abca4^−/−^* mice were significantly higher to those in VAS *Abca4^−/−^* mice (**Figure 3A**, quantified in **Figure 3A’**). At the 7-months of age, 488 nm AF continued to rise in *Rbpr2^−/−^;Abca4^−/−^* mice and were higher than those observed and quantified in VAS *Abca4^−/−^* mice (**Figure 3B**, quantified in **Figure 3B’**). BC supplemented *Abca4^−/−^*mice showed significantly lower fundus AF, compared to both *Rbpr2^−/−^;Abca4^−/−^*and *Abca4^−/−^* mice fed a VAS diet at the 4-month time point (**Figures 3A and 3A’**), and this observation remained consistent at the 7-month-of age timepoint (**Figures 3B and 3B’**). Thus, BC supplementation and the presence of the *Rbpr2* gene attenuated AF lipofuscin accumulation in eyes of *Abca4^−/−^* mice.

### 3.9 Qualitative Optical Coherence Tomography findings in RPE of *Abca4^−/−^* mice

We next conducted a qualitative analysis of retinal pigmented epithelium (RPE) structure in various mice cohorts from above using Optical Coherence Tomography (OCT). In all mice groups a reliable evaluation of RPE skip lesions, thickening, and atrophy was possible. At the 7-month of age, BC supplemented WT and *Abca4^−/−^* mice showed normal/ unremarkable RPE structure (**Figure 4**). Conversely, and at the 7-month of age time-point, *Abca4^−/−^* and *Rbpr2^−/−^;Abca4^−/−^*mice that were fed VAS diets, showed RPE skip lesions (indicated by **\*** in **Figure 4**) and RPE aggregates and thickening (indicated by arrows in **Figure 4**), compared to BC supplemented *Abca4^−/−^* mice. Thus, long-term dietary BC supplementation of *Abca4^−/−^* mice not only attenuated ocular AF lipofuscin accumulation, but also subsequently improved RPE structure, which was dependent on the presence of the *Rbpr2* gene.

**Figure 4:**
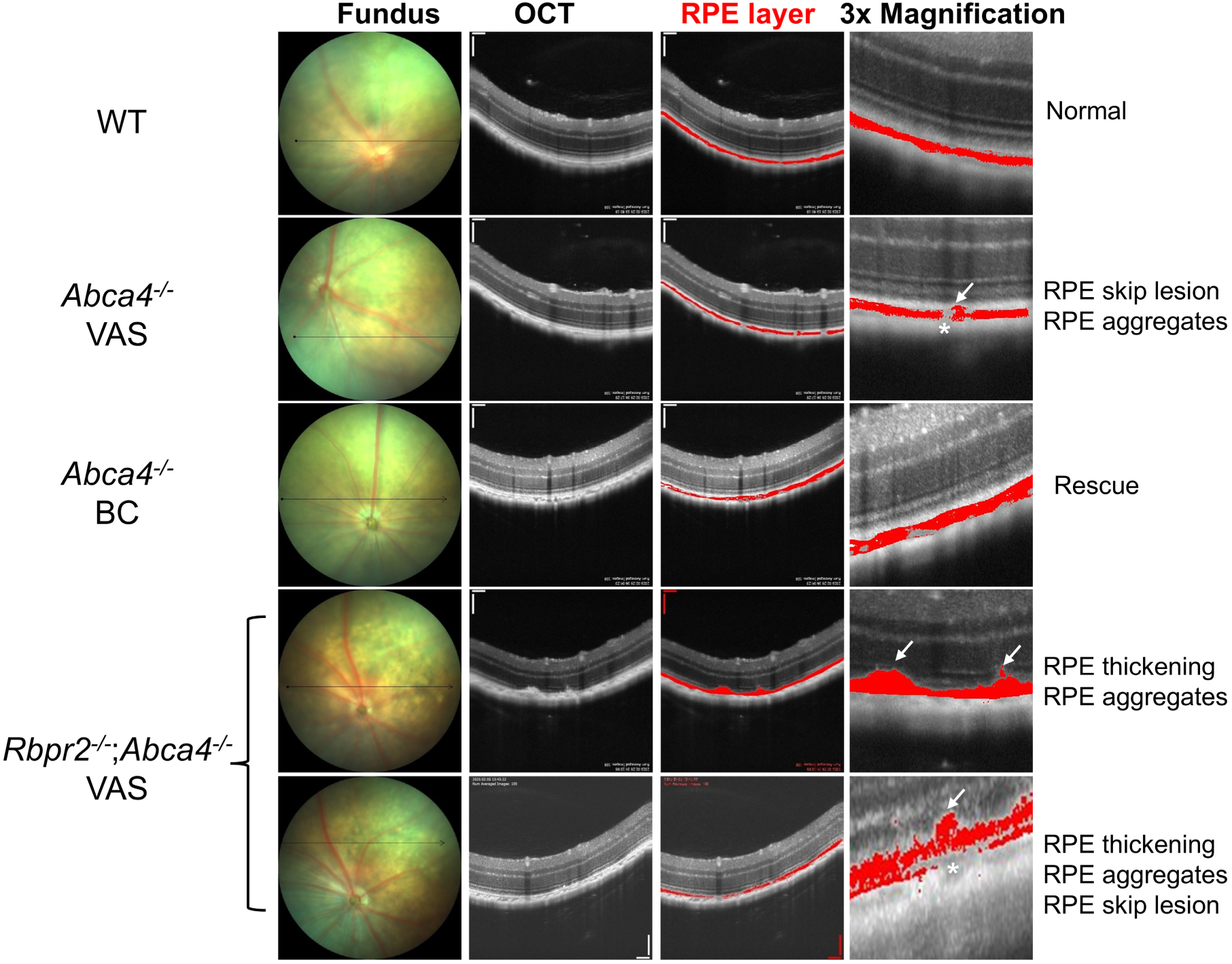
Long-term dietary β-carotene supplementation of *Abca4^−/−^*mice rescues RPE phenotypes. Optical coherence tomography (OCT) was used to evaluate RPE skip lesions, thickening, and atrophy in various cohorts of mice fed either a β-carotene (BC) supplemented or a diet devoid of any β-carotene (VAS diet), at 7-months of age. n=4-6 mice per cohort. VAS, vitamin A sufficient diet; BC, β-carotene supplemented diet. *, RPE skip lesions; arrows, RPE thickening and RPE aggregates.

### 3.10 β-carotene supplemented *Abca4^−/−^* mice show a significant improvement of rod and cone photoreceptor cell function

We then measured rod and cone photoreceptor cell responses by scotopic and photopic flash ERG full-field potentials in the various mice cohorts. After 3– and 6-months of BC supplementation in *Abca4^−/−^*mice, scotopic and photopic ERG responses were measured relative to WT, *Abca4^−/−^*, and *Rbpr2^−/−^;Abca4^−/−^* mice fed a VAS diet of the same age (4– and 7-months-old). We found no significant *a*-wave or *b*-wave differences relative to age-matched WT and *Abca4^−/−^* mice under any feeding conditions in either mouse cohort at 4-months of age (**Supplementary Figures S5A and S5B**). However, at 7-months of age, BC supplemented *Abca4^−/−^* mice showed a significant improvement of rod photoreceptor cell function, compared to either *Abca4^−/−^* or double knockout *Rbpr2^−/−^;Abca4^−/−^* mice fed a VAS diet (**Figures 5A-5H, Supplementary Figure S5C**). Comparison of *b*-wave amplitudes among mice cohorts at the 7-month of age, showed no significant differences between BC supplemented and VAS fed *Abca4^−/−^*mice (**Supplementary Figures S5D, S6A, and S6B**). However, double knockout *Rbpr2^−/−^;Abca4^−/−^* mice fed a VAS diet showed lower *b*-wave amplitudes compared to *Abca4^−/−^* mice fed a VAS or BC supplemented diet (**Supplementary Figures S6C-S6F**).

**Figure 5:**
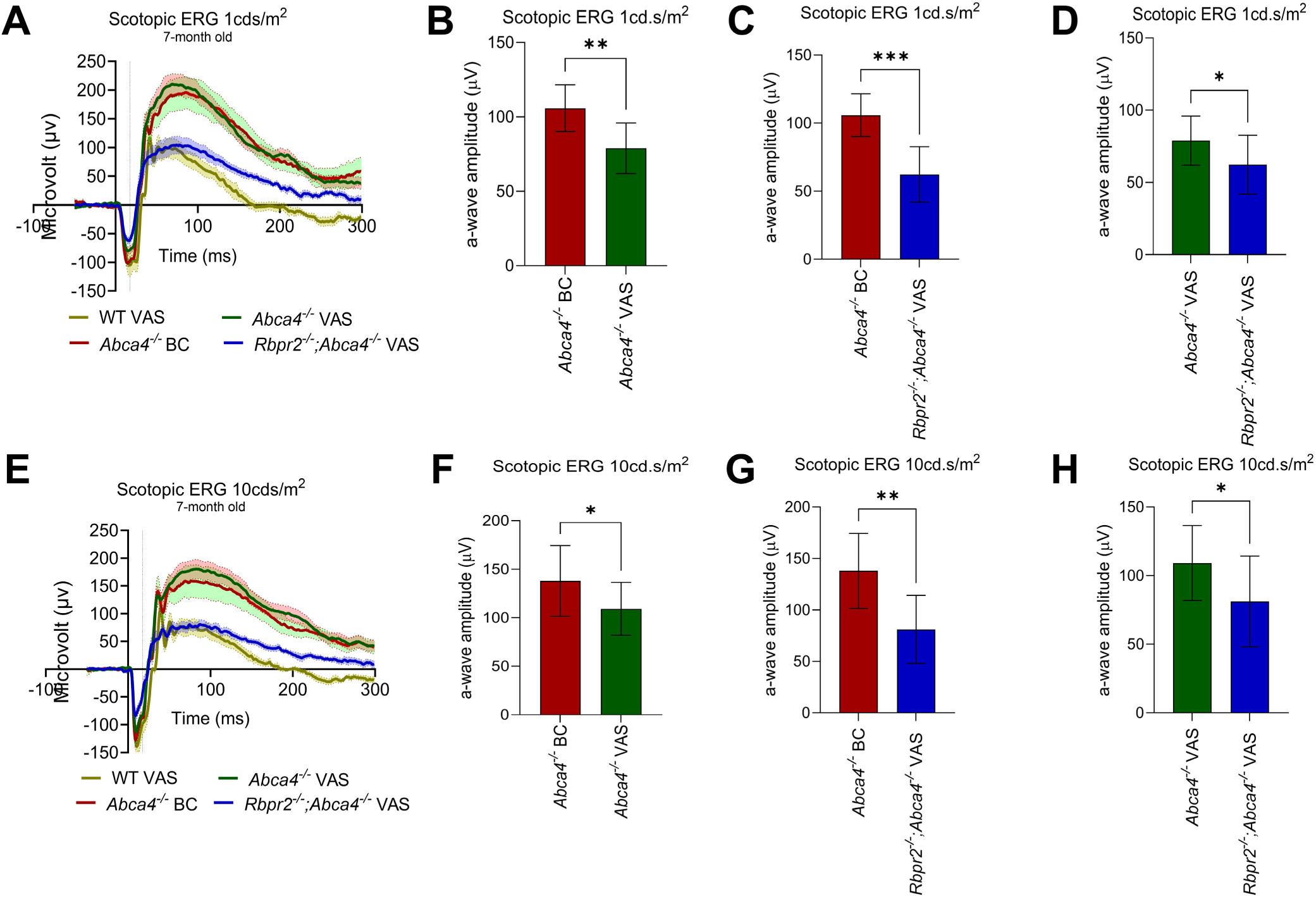
Long-term dietary β-carotene supplementation of *Abca4^−/−^*mice improves rod photoreceptor cell function. Dark-adapted scotopic electroretinographic (ERG) *a*-wave analysis in mice at 7-months of age. (**A-D**) shows scotopic ERG recordings at 1 cd.s/m^2^ and (**E-H**) at 10 cd.s/m^2^. n=4-6 mice per cohort. VAS, vitamin A sufficient diet; BC, β-carotene supplemented diet. Student t-test; *P<0.05; **P<0.01; ***P<0.005.

We next determined ERG responses in light-adapted mice fed either the BC supplemented or VAS diet under different light source color (green, red, white, and UV/blue). Under photopic light conditions, ERG responses of *Abca4^−/−^* mice under green, white, and UV color sources were significantly diminished at the 7-month timepoint, while red light source ERG responses were not changed, when compared with BC supplemented *Abca4^−/−^* mice (**Supplementary Figures S7A-S7D**). Light-adapted ERG responses for blue (**Supplementary Figures S7A, S7A’, and S7A’’**), green (**Supplementary Figures S7B, S7B’, and S7B’’**), and white (**Supplementary Figures S7C, S7C’, and S7C’’**) light intensity improved in BC supplemented *Abca4^−/−^*mice at the 7-month timepoint, but remained significantly diminished under red-light exposure (**Supplementary Figure S7D**). Thus, BC supplemented *Abca4^−/−^* mice showed an overall improvement of rod and cone photoreceptor cell function, compared to *Abca4^−/−^* mice fed a VAS diet or double knockout *Rbpr2^−/−^;Abca4^−/−^* mice.

### 3.11 β-carotene supplementation of *Abca4^−/−^* mice improves RPE function

A key pathological feature of STGD1 is the progressive accumulation of fluorescent lipofuscin/A2E pigments in the retinal pigmented epithelium (RPE), which could potentially affect RPE function^30^ and be causative of photoreceptor dysfunction observed above. We therefore used *c*-wave ERGs to test for RPE function in the various mice cohorts. At 7-months of age BC supplemented *Abca4^−/−^*mice showed an improvement of RPE function, compared to either *Abca4^−/−^*or double knockout *Rbpr2^−/−^;Abca4^−/−^* mice fed a VAS diet (**Figures 6A-6C**). Additionally, double knockout *Rbpr2^−/−^;Abca4^−/−^*mice showed an even more severe decrease in RPE function, compared to VAS *Abca4^−/−^* mice (**Figure 6D**).

**Figure 6:**
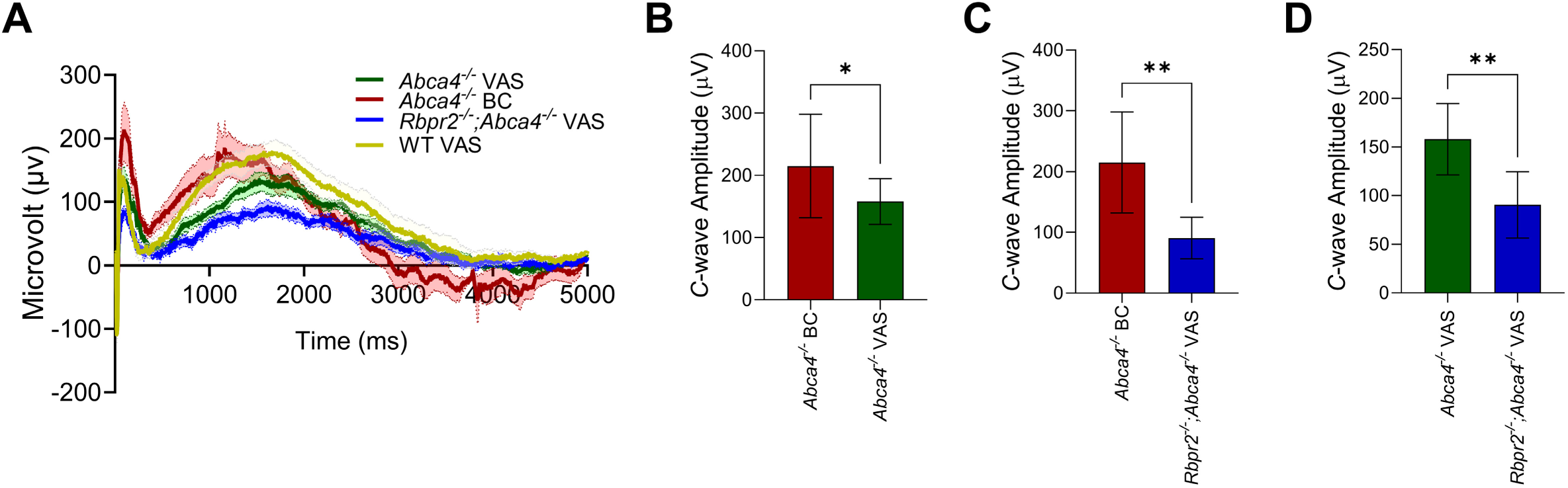
Long-term dietary β-carotene supplementation of *Abca4^−/−^*mice improves RPE cell function. (**A-D**) Dark adapted *c*-wave analysis for RPE cell function in 7-month-old mice among different genotypes and dietary conditions. n=4-6 mice per cohort. VAS, vitamin A sufficient diet; BC, β-carotene supplemented diet. Student t-test; *P<0.05; **P<0.01.

### 3.12 Quantification of A2E isomers in mice eyes by High Performance Liquid Chromatography

Since loss of *ABCA4* has been associated with accumulation of lipofuscin pigments in the eye, we quantified A2E, a bisretinoid component of lipofuscin, by integrating HPLC peaks and normalizing to a calibration curve obtained from an A2E and iso-A2E standard **(Supplementary Figure S8)**. HPLC analysis revealed that A2E and iso-A2E isomer concentrations in 7-month-old BC supplemented *Abca4^−/−^* mice were significantly lower, compared to those fed a VAS diet (**Figures 7A-7E**), which also collated with the fundus AF images observed in these mice cohorts (**Figure 3B**). Comparison of A2E and iso-A2E isomer concentrations in 7-month-old *Abca4^−/−^* and double knockout *Rbpr2^−/−^;Abca4^−/−^*mice fed a VAS diet, showed no significant differences (P<0.0601; **Figure 7B**).

**Figure 7:**
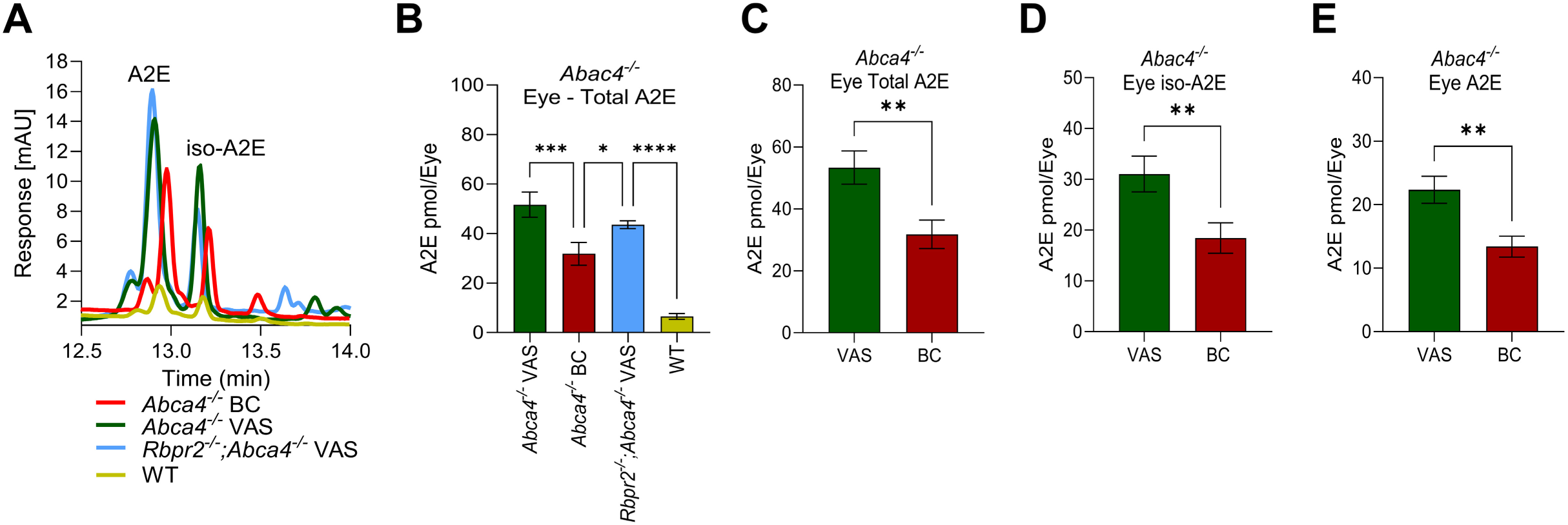
HPLC quantification of A2E and iso-A2E isomers in mice eyes. (**A**) HPLC traces of A2E and iso-A2E isomers in *Abca4^−/−^* and *Rbpr2^−/−^;Abca4^−/−^* mice eyes, fed a VAS or BC supplement diet, at 7-months of age. (**B**) Total ocular A2E concentrations shown in pmole/eye among various mice cohorts. (**C**) Total A2E, (**D**) iso-A2E, and (**E**) A2E, in eyes of *Abca4^−/−^*mice fed either a VAS or BC supplemented diet. n=4-6 mice per cohort. VAS, vitamin A sufficient diet; BC, β-carotene supplemented diet. One way ANOVA or Student t-test; *P<0.05; **P<0.01; ***P<0.005; ****P<0.001.

### 3.13 Fluorescence Microscopy of Mice Retinal Sections for Lipofuscin Accumulation

We next used retinal sections and fluorescence microscopy to visualize A2E/lipofuscin accumulation in the RPE of various mice cohorts, fed either a VAS or BC supplemented diet. This analysis revealed that A2E/ lipofuscin fluorescence in 7-month-old *Abca4^−/−^* mice supplemented with dietary BC were significantly lower to those on a VAS diet (**Figure 8A**). This observation also collated with the fundus AF images observed in these mice cohorts (**Figure 3B**) and with the HPLC quantification of A2E isomers (**Figure 7**). Comparison of A2E florescence among mice cohorts, showed that double knockout *Rbpr2^−/−^;Abca4^−/−^*mice displayed higher A2E/lipofuscin fluorescence intensity, compared to VAS *Abca4^−/−^* mice (**Figure 8B**). Thus, this data supports the hypothesis that the systemic vitamin A receptor (*Rbpr2*) in synergy with a carotenoid (β-carotene) supplemented diet, plays a beneficial role in attenuating lipofuscin/A2E accumulation, which leads to an improvement of photoreceptor and RPE function in *Abca4^−/−^*mice.

**Figure 8:**
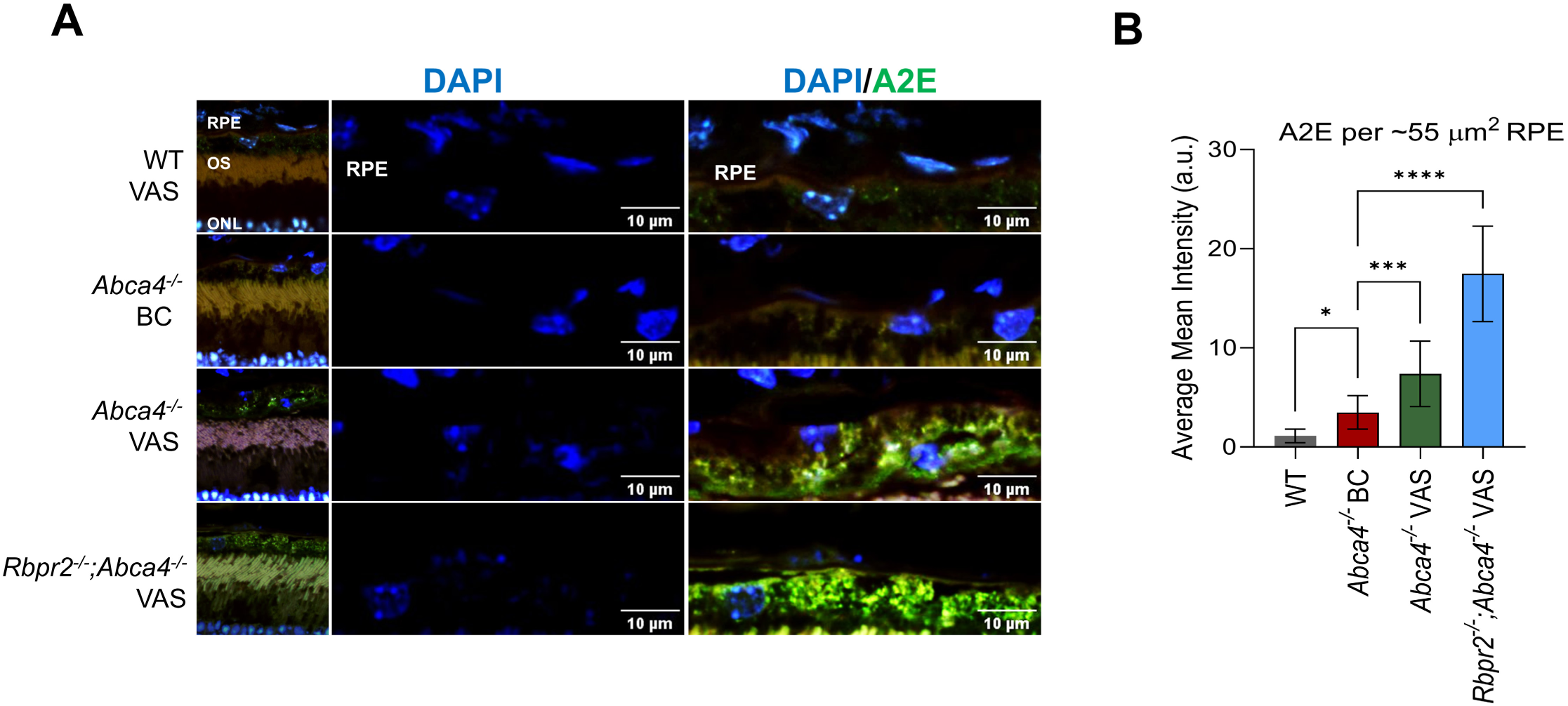
Fluorescent microscopy detects autofluorescent lipofuscin/ A2E granules in RPE of mice eyes. Autofluorescent lipofuscin/ A2E granules in RPE of mice fed either a VAS or BC supplemented diet, at 7-months of age. (**A**) Two channels were used with 405 nm excitation and emission of 570-616 nm and 607-850 nm to detect A2E fluorescence in RPE/retinas from various mice cohorts. (**B**) Quantification of autofluorescent lipofuscin/ A2E granules in RPE. n=4-6 mice per cohort. VAS, vitamin A sufficient diet; BC, β-carotene supplemented diet. One way ANOVA, *P<0.05; ***P<0.005; *****P<0.001.

### 3.14 β-carotene supplemented *Abca4^−/−^* mice show rescue of cone photoreceptor phenotypes

Since lipofuscin/A2E accumulation in the absence of *ABCA4* has been linked to macular degeneration, which can specifically affect integrity of cone photoreceptors, we next performed immunohistology analysis of retinas in the different mice cohorts for cone photoreceptor morphology using R/G cone opsin staining. This analysis showed that VAS *Abca4^−/−^* mice and double knockout *Rbpr2^−/−^;Abca4^−/−^*mice at 7-months of age had shorter and misshaped cone outer segments (OS), compared to WT mice at the same age (**Figure 9A**). Interestingly, double knockout *Rbpr2^−/−^;Abca4^−/−^*mice showed an even shorter cone photoreceptor OS phenotype, compared to *Abca4^−/−^*mice (**Figure 9**, cone length quantified in **Figure 9B**). Conversely, BC supplemented *Abca4^−/−^*mice showed a rescue of cone OS phenotypes, indicating that long-term BC supplemented diet and presence of the vitamin A receptor, *Rbpr2*, can protect against cone photoreceptor OS phenotypes in *Abca4^−/−^* mice (**Figures 9A and 9B**).

**Figure 9:**
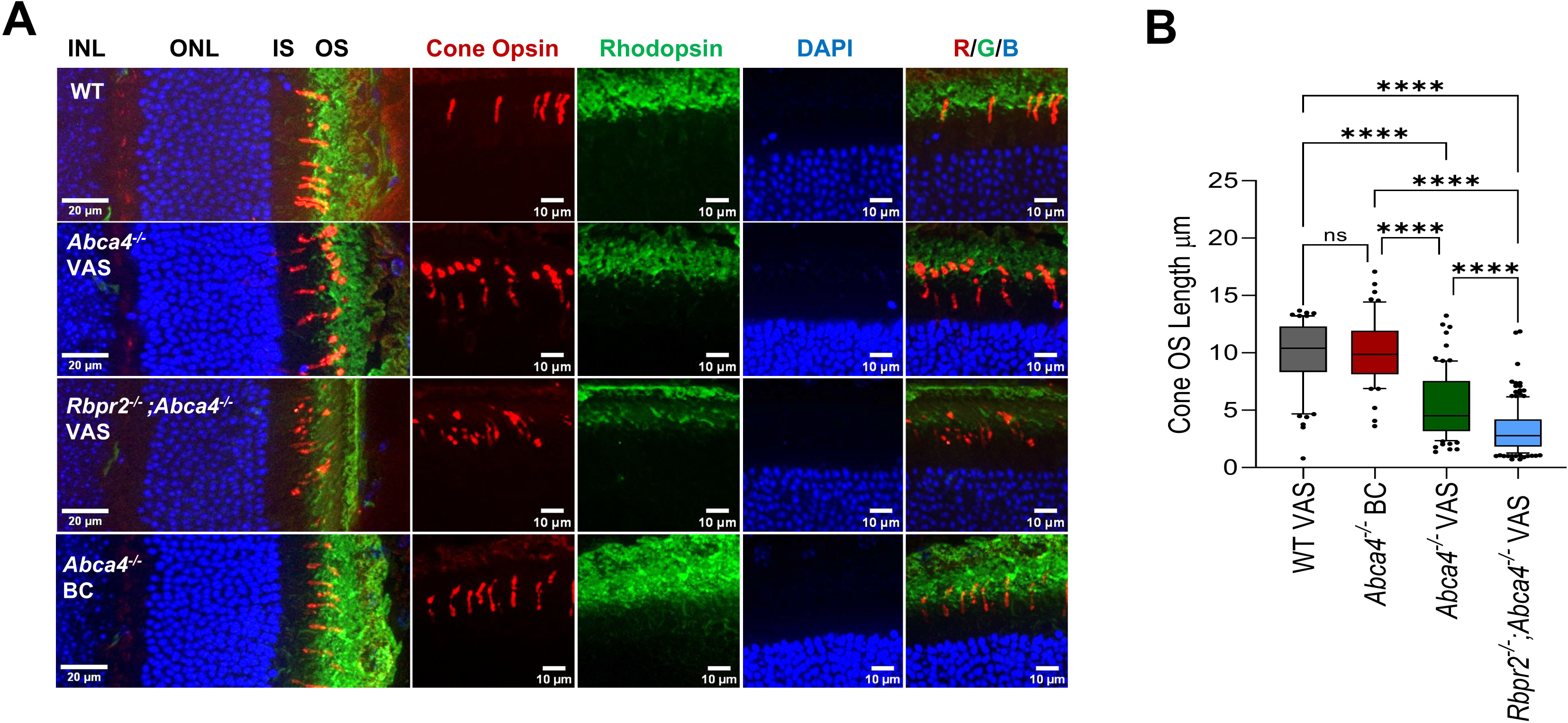
Long-term dietary β-carotene supplementation of *Abca4^−/−^*mice rescues cone photoreceptor phenotypes. (**A**) Retinal sections from various mouse cohorts at 7-months of age were subjected to immunohistochemistry using 1D4 (rhodopsin; green) and R/G cone opsin (red) antibodies. DAPI (blue) in the mounting solution was used to stain nuclei. (**B**) Quantification of cone photoreceptor length from various mouse cohorts. n=3-5 retinal sections from n=3-6 mice eyes per cohort. VAS, vitamin A sufficient diet; BC, β-carotene supplemented diet. One way ANOVA, ****P<0.001.

## 4. DISCUSSION

Here we identify a synergistic role between long-term dietary β-carotene (BC) supplementation and presence of the non-ocular vitamin A receptor *Rbpr2* gene as a therapeutic modality in attenuating cytotoxic lipofuscin/A2E accumulation in *Abca4^−/−^* mice, a well-established mouse model of Stargardt disease/STGD1 (**Figure 10**). Since serum all-*trans*-retinol (ROL) serves as the first step precursor for ocular all-*trans*-retinal synthesis, a component of cytotoxic A2E, we explored in vivo mechanisms that could regulate serum ROL bound RBP4 (RBP4-ROL) concentration in *Abca4^−/−^*mice. We identify multiple RAR and RXR sites on the vitamin A receptor *Rbpr2* gene promotor, which can be induced in vitro and in vivo by BC metabolites, all-*trans*-retinoic acid (RA), 9-*cis*-retinoic acid (9cRA), and 13-cis-retinoic acid (13cRA). In vivo, this mechanism enhanced hepatic retinoic acid production and significantly induced *Rbpr2* mRNA expression, reducing serum RBP4 levels in WT and *Abca4^−/−^* mice. Subsequently, long-term dietary BC supplementation of *Abca4^−/−^* mice, reduced fundus auto fluorescence (AF) levels, decreased ocular A2E and iso-A2E isomer accumulation in the RPE, which led to an improvement of rod and cone photoreceptor and RPE cell function. Conversely, neither *Abca4^−/−^*or double knockout *Rbpr2^−/−^;Abca4^−/−^* mice fed a diet devoid of BC, showed such a rescue of ocular phenotypes. Thus, we identified a novel mechanism, where long-term dietary BC supplementation and subsequent metabolism to RA isomers influences activity of the systemic vitamin A receptor *Rbpr2,* which in turn regulates serum RBP4-ROL precursors required for cytotoxic A2E production in *Abca4^−/−^* mice. This mechanism we hypothesize could benefit STGD1 patients with *ABCA4* gene mutations.

**Figure 10:**
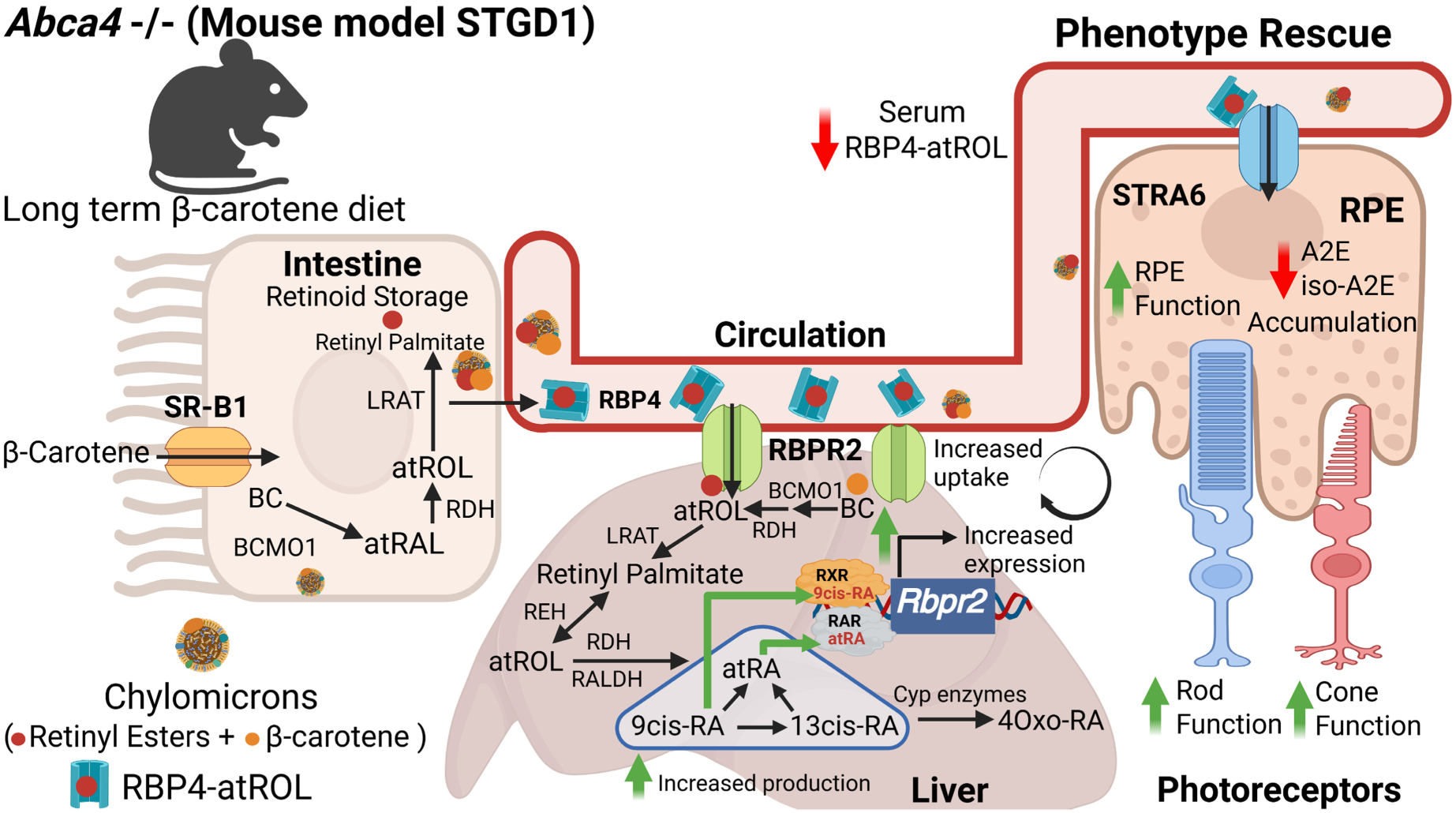
Genetics and the Diet attenuate ocular Lipofuscin/A2E production and accumulation and improves visual and RPE function in *Abca4^−/−^* mice. *Abca4^−/−^*mice were fed a β-carotene (BC) supplemented or vitamin A sufficient (VAS) diet for up to 7-months of age. BC supplemented *Abca4^−/−^* mice showed a higher production of retinoic acid isomers (atRA, 9*cis*RA, and 13*cis*RA) and BC concentrations in the liver, which led to an induction of hepatic *Rbpr2* gene activity via RAR and RXR regulatory elements on its promoter. This mechanism increased reuptake of serum ROL bound RBP4 by *Rbpr2*, in turn reducing serum RBP4 concentrations. In the long-term, this mechanism reduced ocular A2E and iso-A2E production and accumulation, overall improving rod and cone photoreceptor, and RPE function in *Abca4^−/−^* mice fed BC supplemented diets. BC, β-carotene; VAS, vitamin A sufficient; atRA, all-*trans*-retinoic acid; 9*cis*RA, 9-*cis*-retinoic acid; 13*cis*RA, 13-*cis*-retinoic acid; BCMO1, β-carotene monooxygenase; LRAT, Lecithin Retinol Acyltransferase; atROL, all-*trans*-retinol; RDH, retinol dehydrogenase; RALDH, retinal dehydrogenase; RAR, retinoic acid receptor; RXR, retinoid X receptor; 4Oxo-RA, 4-oxo-retinoic acid; RBP4, retinol binding protein 4; RBPR2, retinol binding protein 4 receptor 2; SR-B1, Scavenger receptor class B receptor 1; STRA6, stimulated by retinoic acid 6; STGD1, Stargar<u>d</u>t D<u>is</u>ease; *<u>A</u>bca4*, ATP-binding cassette subfamily <u>A</u> member <u>4</u> transporter. Schematic figure created in BioRender (BioRender.com/a90mdtq).

First, we confirmed and expanded on the previously proposed transcriptional regulation of the systemic vitamin A receptor *Rbpr2*^18^. We and others have shown that *Rbpr2* is highly expressed in the murine liver, intestine, and adipose tissue, followed by moderate to lower expression in other non-ocular tissues^10,18,20,31^. Because these tissues mediate uptake, storage, and distribution of all-*trans*-retinol (ROL) to peripheral tissues including the eye, these observations suggest that RBPR2 could play a physiological role in whole-body ROL/retinoid homeostasis^10,18,20^. Additionally, the Graham laboratory had previously suggested negative regulation of the murine *Rbpr2* gene promotor activity via retinoic acid response elements (RARE) on its promotor, which could be relevant in regulating liver ROL stores^18^. Our detailed analysis of the *Rbpr2* gene promotor region revealed multiple RAR and RXR sites, which were transcriptionally regulated by retinoids. We also confirmed previous observations that exogenous all-*trans*-ROL can downregulate *Rbpr2* mRNA expression and further discovered that exogenous BC metabolites induce *Rbpr2* mRNA expression in cultured cells. The latter observation was then confirmed in vivo in BC fed WT mice. The mechanism for induction of hepatic *Rbpr2* gene activity in BC supplemented WT and *Abca4^-/^*^-^ mice we attributed to significantly higher production of hepatic RA metabolites (atRA, 9cRA, and 13cRA) derived from BC metabolism, compared to VAS fed animals. Interestingly and more recently, hepatic *Rbpr2* mRNA expression was also found to be significantly induced in BC fed animals of a different genetic background, which led to increased accumulation of hepatic retinoids, further indicating that induction of hepatic *Rbpr2* gene expression can lead to increased re-absorption of circulatory ROL from RBP4^32^. Thus, based on our findings here and those previously reported by other investigators, exogenous and dietary BC metabolites induces *Rbpr2* gene activity via RARE elements on its promoter region, which plays a physiological role in modulating circulatory ROL bound RBP4 levels (**Figure 10**).

Stargardt disease (STGD1) is the phenotype most caused by *ABCA4* gene mutations. A key pathological feature of STGD1 is the buildup of fundus autofluorescence (AF) lipofuscin/bisretinoid A2E pigments in the retinal pigmented epithelium (RPE), a feature previously reported in *Abca4^−/−^*mice. We note that the effects of genetic deletion of the vitamin A receptor, *Rbpr2*, on the *Abca4^−/−^* background (double knockout *Rbpr2^−/−^;Abca4^−/−^* mice) was not only additive to the *Abca4^−/−^*mice ocular phenotypes, but was also causative of earlier appearance and detection of fundus AF lipofuscin. This observation was made by comparing the fundus AF images from *Abca4^−/−^* and *Rbpr2^−/−^;Abca4^−/−^*mice fed a VAS diet, at the earlier 4-month of age and then at 7-month of age time-point. Specifically at the 4-month of age, fundus AF levels in *Rbpr2^−/−^;Abca4^−/−^*mice fed a diet devoid of BC (VAS diet) were observed to be significantly higher, to those noted in *Abca4^−/−^* mice also fed the same VAS diet, indicating that loss of the *Rbpr2* receptor in these mice was additive to the retinal phenotypes. Among the explanations for this observation could be the unregulated and higher levels of serum RBP4-ROL in VAS *Rbpr2^−/−^;Abca4^−/−^* mice, which could have provided additional precursors for accelerating ocular A2E accumulation. Conversely, *Abca4^−/−^* mice fed a BC diet showed increased hepatic *Rbpr2* gene expression and decreased serum RBP4 levels, which was reflected in reduced fundus AF levels, indicating that both genetics (*Rbpr2* gene presence) and the diet (BC) acts as a therapeutic modality, in modulating serum RBP4-ROL levels in these mice. In fact, long-term BC supplementation, but not VAS diets, in *Abca4^−/−^*mice, showed a significant reduction in A2E/lipofuscin accumulation in the RPE of these animals, which improved photoreceptor and RPE cell function.

Our study has certain limitations, which can be addressed by generating novel mouse models in the future. The reductions in serum RBP4-ROL in BC supplemented WT and *Abca4^−/−^*mice, might be a due to a combination of different mechanisms, either due to increased hepatic re-uptake of circulatory ROL from RBP4 via BC metabolite production and induction of *Rbpr2* gene expression (observed herein), or due to increased storage of vitamin A (as retinyl esters) in the intestine, or due to increased efflux of ROL from the intestine by RBPR2 (**Figure 10**). Since, RBPR2 has been shown by us and others to be highly expressed in the murine intestine (presumably basolateral) and in the liver, we are currently establishing tissue-specific liver (*Rbpr2^−/−^;Alb-Cre+*) and intestine (*Rbpr2^−/−^;Villin-Cre+*) mice to answer this question.

Importantly, since vitamin A content in western diets predominantly consists of high-levels of preformed retinoids (>75%), it can be hypothesized that the un-regulated and excess ocular uptake of circulatory preformed retinoids, could contribute to accelerated A2E/lipofuscin biogenesis in humans^33,34^. Based on these observations, we propose that dietary *β*-carotene derived RA regulation of systemic *Rbpr2* receptor activity might serve to limit excess circulatory and ocular ROL precursor availability. This regulatory mechanism via vitamin A transporters could attenuate accelerated toxic ocular A2E biosynthesis in patients with *ABCA4* mutations, thus preserving visual function in STGD1 patients on long-term carotenoid sufficient diets^35^.

In summary, we have shown that long-term dietary BC supplementation in *Abca4^−/−^* mice, reduced serum RBP4 levels via *Rbpr2* gene activity, which translated to robust reduction of cytotoxic A2E and iso-A2E accumulation in the RPE, which in the long-term improved photoreceptor and RPE function (**Figure 10**). Importantly, these positive effects of BC diet in *Abca4^−/−^*mice, were dependent on the presence of the non-ocular vitamin A receptor *Rbpr2*. Thus, our approach could not only benefit STGD1 patients with *ABCA4* gene mutations, but in the future could also be explored as a therapeutic modality in other mouse models of macular degeneration, such as *MnSod^−/−^*, *Abcr^−/−^*, and *Elvol4^−/−^* mice^36–41^, where A2E/lipofuscin pigmented granule accumulation is the major cause of RPE and photoreceptor cell degeneration in these mutant mouse models of STGD1.

## AUTHOR CONTRIBUTIONS

Conceptualization, G.P.L.; methodology, G.P.L., R.R., M.L., H.R.; software, R.R.; validation, R.R., G.P.L., M.L.; reagents, G.P.L., S.M.; formal analysis, R.R., G.P.L., D.Y., H.R., M.L.; investigation, R.R., G.P.L., M.L.; resources, G.P.L., S.M., H.R.; data curation, R.R., G.P.L., M.L.; writing-original draft preparation, G.P.L.; manuscript writing, review, and editing, R.R., G.P.L., M.L., D.Y., S.M.; supervision, G.P.L.; project administration, G.P.L.; funding acquisition, G.P.L. All authors have read and agreed to the published version of the manuscript.

## Abbreviations

ABCA4: ATP-binding cassette transporter subfamily A4
A2E: *N*-retinyl-*N*-retinylidene ethanolamine (pyridinium bisretinoid)
BC: β-carotene
ERG: Electroretinogram
RAR: retinoic acid receptor
RARE: retinoic acid response element
RBPR2: retinol binding protein 4 receptor 2
RBP4: retinol binding protein 4
ROL: all-*trans*-retinol
RA: all-*trans*-retinoic acid
RPE: retinal pigmented epithelium
STGD1: Stargardt Disease Type 1
HPLC: High Performance Liquid Chromatography

## ACKNOWLEDGMENTS

This work was supported by NIH-NEI grants (EY030889 and 3R01EY030889-03S1 G.P.L. and EY033328 to S.W.M.) and in part by the University of Minnesota start-up funds to G.P.L.

## DATA AVAILABILITY STATEMENT

Data sharing is not applicable to this article, as no datasets were generated or analyzed in this study. Reagents are available from the corresponding author upon reasonable request.

## REFERENCES

(1) Sahel, J.-A.; Marazova, K.; Audo, I. Clinical Characteristics and Current Therapies for Inherited Retinal Degenerations. Cold Spring Harb. Perspect. Med. 2015, 5 (2), a017111. 10.1101/cshperspect.a017111.

(2) Michaelides, M.; Hunt, D. M.; Moore, A. T. The Cone Dysfunction Syndromes. Br. J. Ophthalmol. 2004, 88 (2), 291–297. 10.1136/bjo.2003.027102.

(3) Fujinami, K.; Zernant, J.; Chana, R. K.; Wright, G. A.; Tsunoda, K.; Ozawa, Y.; Tsubota, K.; Robson, A. G.; Holder, G. E.; Allikmets, R.; Michaelides, M.; Moore, A. T. Clinical and Molecular Characteristics of Childhood-Onset Stargardt Disease. Ophthalmology 2015, 122 (2), 326–334. 10.1016/j.ophtha.2014.08.012.

(4) Michaelides, M.; Hunt, D. M.; Moore, A. T. The Genetics of Inherited Macular Dystrophies. J. Med. Genet. 2003, 40 (9), 641–650. 10.1136/jmg.40.9.641.

(5) Fujinami, K.; Lois, N.; Davidson, A. E.; Mackay, D. S.; Hogg, C. R.; Stone, E. M.; Tsunoda, K.; Tsubota, K.; Bunce, C.; Robson, A. G.; Moore, A. T.; Webster, A. R.; Holder, G. E.; Michaelides, M. A Longitudinal Study of Stargardt Disease: Clinical and Electrophysiologic Assessment, Progression, and Genotype Correlations. Am. J. Ophthalmol. 2013, 155 (6), 1075–1088.e13. 10.1016/j.ajo.2013.01.018.

(6) McBain, V. A.; Townend, J.; Lois, N. Progression of Retinal Pigment Epithelial Atrophy in Stargardt Disease. Am. J. Ophthalmol. 2012, 154 (1), 146–154. 10.1016/j.ajo.2012.01.019.

(7) Fishman, G. A.; Stone, E. M.; Grover, S.; Derlacki, D. J.; Haines, H. L.; Hockey, R. R. Variation of Clinical Expression in Patients With Stargardt Dystrophy and Sequence Variations in the ABCRGene. Arch. Ophthalmol. 1999, 117 (4), 504–510. 10.1001/archopht.117.4.504.

(8) Lambertus, S.; Huet, R. A. C. van; Bax, N. M.; Hoefsloot, L. H.; Cremers, F. P. M.; Boon, C. J. F.; Klevering, B. J.; Hoyng, C. B. Early-Onset Stargardt Disease: Phenotypic and Genotypic Characteristics. Ophthalmology 2015, 122 (2), 335–344. 10.1016/j.ophtha.2014.08.032.

(9) Vance, J. E. Thematic Review Series: Glycerolipids. Phosphatidylserine and Phosphatidylethanolamine in Mammalian Cells: Two Metabolically Related Aminophospholipids. J. Lipid Res. 2008, 49 (7), 1377–1387. 10.1194/jlr.R700020-JLR200.

(10) Martin Ask, N.; Leung, M.; Radhakrishnan, R.; Lobo, G. P. Vitamin A Transporters in Visual Function: A Mini Review on Membrane Receptors for Dietary Vitamin A Uptake, Storage, and Transport to the Eye. Nutrients 2021, 13 (11), 3987. 10.3390/nu13113987.

(11) Leung, M.; Steinman, J.; Li, D.; Lor, A.; Gruesen, A.; Sadah, A.; van Kuijk, F. J.; Montezuma, S. R.; Kondkar, A. A.; Radhakrishnan, R.; Lobo, G. P. The Logistical Backbone of Photoreceptor Cell Function: Complementary Mechanisms of Dietary Vitamin A Receptors and Rhodopsin Transporters. Int. J. Mol. Sci. 2024, 25 (8), 4278. 10.3390/ijms25084278.

(12) Scortecci, J. F.; Molday, L. L.; Curtis, S. B.; Garces, F. A.; Panwar, P.; Van Petegem, F.; Molday, R. S. Cryo-EM Structures of the ABCA4 Importer Reveal Mechanisms Underlying Substrate Binding and Stargardt Disease. Nat. Commun. 2021, 12 (1), 5902. 10.1038/s41467-021-26161-7.

(13) Kiser, P. D.; Golczak, M.; Palczewski, K. Chemistry of the Retinoid (Visual) Cycle. Chem. Rev. 2014, 114 (1), 194–232. 10.1021/cr400107q.

(14) Sparrow, J. R.; Gregory-Roberts, E.; Yamamoto, K.; Blonska, A.; Ghosh, S. K.; Ueda, K.; Zhou, J. The Bisretinoids of Retinal Pigment Epithelium. Prog. Retin. Eye Res. 2012, 31 (2), 121–135. 10.1016/j.preteyeres.2011.12.001.

(15) Sparrow, J. R.; Fishkin, N.; Zhou, J.; Cai, B.; Jang, Y. P.; Krane, S.; Itagaki, Y.; Nakanishi, K. A2E, a Byproduct of the Visual Cycle. Vision Res. 2003, 43 (28), 2983–2990. 10.1016/S0042-6989(03)00475-9.

(16) Sparrow, J. R.; Nakanishi, K.; Parish, C. A. The Lipofuscin Fluorophore A2E Mediates Blue Light–Induced Damage to Retinal Pigmented Epithelial Cells. Invest. Ophthalmol. Vis. Sci. 2000, 41 (7), 1981–1989.

(17) Ben-Shabat, S.; Parish, C. A.; Vollmer, H. R.; Itagaki, Y.; Fishkin, N.; Nakanishi, K.; Sparrow, J. R. Biosynthetic Studies of A2E, a Major Fluorophore of Retinal Pigment Epithelial Lipofuscin*. J. Biol. Chem. 2002, 277 (9), 7183–7190. 10.1074/jbc.M108981200.

(18) Alapatt, P.; Guo, F.; Komanetsky, S. M.; Wang, S.; Cai, J.; Sargsyan, A.; Rodríguez Díaz, E.; Bacon, B. T.; Aryal, P.; Graham, T. E. Liver Retinol Transporter and Receptor for Serum Retinol-Binding Protein (RBP4). J. Biol. Chem. 2013, 288 (2), 1250–1265. 10.1074/jbc.M112.369132.

(19) Lobo, G. P.; Pauer, G.; Lipschutz, J. H.; Hagstrom, S. A. The Retinol-Binding Protein Receptor 2 (Rbpr2) Is Required for Photoreceptor Survival and Visual Function in the Zebrafish. In Retinal Degenerative Diseases; Ash, J. D., Anderson, R. E., LaVail, M. M., Bowes Rickman, C., Hollyfield, J. G., Grimm, C., Eds.; Advances in Experimental Medicine and Biology; Springer International Publishing: Cham, 2018; pp 569–576. 10.1007/978-3-319-75402-4_69.

(20) Radhakrishnan, R.; Leung, M.; Lor, A.; More, S.; Lobo, G. P. Loss of the Vitamin A Receptor RBPR2 in Mice Disrupts Whole-Body Retinoid Homeostasis and the Quantitative Balance Regulating Retinylidene Protein Synthesis. FASEB J. 2025, 39 (5), e70407. 10.1096/fj.202403090R.

(21) Lobo, G. P.; Hessel, S.; Eichinger, A.; Noy, N.; Moise, A. R.; Wyss, A.; Palczewski, K.; von Lintig, J. ISX Is a Retinoic Acid-Sensitive Gatekeeper That Controls Intestinal β,β-Carotene Absorption and Vitamin A Production. FASEB J. 2010, 24 (6), 1656–1666. 10.1096/fj.09-150995.

(22) Lobo, G. P.; Amengual, J.; Baus, D.; Shivdasani, R. A.; Taylor, D.; von Lintig, J. Genetics and Diet Regulate Vitamin A Production via the Homeobox Transcription Factor ISX. J. Biol. Chem. 2013, 288 (13), 9017–9027. 10.1074/jbc.M112.444240.

(23) Leung, M.; Radhakrishnan, R.; Lor, A.; Li, D.; Yochim, D.; More, S.; Lobo, G. P. Quantitative Analysis of Dietary Vitamin A Metabolites in Murine Ocular and Non-Ocular Tissues Using High-Performance Liquid Chromatography. J. Vis. Exp. JoVE 2024, No. 214, e67034. 10.3791/67034.

(24) Sparrow, J. R.; Kim, S. R.; Wu, Y. Experimental Approaches to the Study of A2E, a Bisretinoid Lipofuscin Chromophore of Retinal Pigment Epithelium. Methods Mol. Biol. Clifton NJ 2010, 652, 315–327. 10.1007/978-1-60327-325-1_18.

(25) Zhao, J.; Kim, H. J.; Ueda, K.; Zhang, K.; Montenegro, D.; Dunaief, J. L.; Sparrow, J. R. A Vicious Cycle of Bisretinoid Formation and Oxidation Relevant to Recessive Stargardt Disease. J. Biol. Chem. 2021, 296, 100259. 10.1016/j.jbc.2021.100259.

(26) Sparrow, J. R.; Blonska, A.; Flynn, E.; Duncker, T.; Greenberg, J. P.; Secondi, R.; Ueda, K.; Delori, F. C. Quantitative Fundus Autofluorescence in Mice: Correlation With HPLC Quantitation of RPE Lipofuscin and Measurement of Retina Outer Nuclear Layer Thickness. Invest. Ophthalmol. Vis. Sci. 2013, 54 (4), 2812–2820. 10.1167/iovs.12-11490.

(27) Boyer, N. P.; Higbee, D.; Currin, M. B.; Blakeley, L. R.; Chen, C.; Ablonczy, Z.; Crouch, R. K.; Koutalos, Y. Lipofuscin and *N*-Retinylidene-*N*-Retinylethanolamine (A2E) Accumulate in Retinal Pigment Epithelium in Absence of Light Exposure. J. Biol. Chem. 2012, 287 (26), 22276–22286. 10.1074/jbc.M111.329235.

(28) Moiseyev, G.; Nikolaeva, O.; Chen, Y.; Farjo, K.; Takahashi, Y.; Ma, J. Inhibition of the Visual Cycle by A2E through Direct Interaction with RPE65 and Implications in Stargardt Disease. Proc. Natl. Acad. Sci. 2010, 107 (41), 17551–17556. 10.1073/pnas.1008769107.

(29) Charbel Issa, P.; Barnard, A. R.; Singh, M. S.; Carter, E.; Jiang, Z.; Radu, R. A.; Schraermeyer, U.; MacLaren, R. E. Fundus Autofluorescence in the Abca4(−/−) Mouse Model of Stargardt Disease--Correlation with Accumulation of A2E, Retinal Function, and Histology. Invest. Ophthalmol. Vis. Sci. 2013, 54 (8), 5602–5612. 10.1167/iovs.13-11688.

(30) Guan, Z.; Li, Y.; Jiao, S.; Yeasmin, N.; Rosenfeld, P. J.; Dubovy, S. R.; Lam, B. L.; Wen, R. A2E Distribution in RPE Granules in Human Eyes. Mol. Basel Switz. 2020, 25 (6), 1413. 10.3390/molecules25061413.

(31) Radhakrishnan, R.; Leung, M.; Roehrich, H.; Walterhouse, S.; Kondkar, A. A.; Fitzgibbon, W.; Biswal, M. R.; Lobo, G. P. Mice Lacking the Systemic Vitamin A Receptor RBPR2 Show Decreased Ocular Retinoids and Loss of Visual Function. Nutrients 2022, 14 (12), 2371. 10.3390/nu14122371.

(32) Moon, J.; Ramkumar, S.; von Lintig, J. Genetic Dissection in Mice Reveals a Dynamic Crosstalk between the Delivery Pathways of Vitamin A. J. Lipid Res. 2022, 63 (6), 100215. 10.1016/j.jlr.2022.100215.

(33) Petrukhin, K. Pharmacological Inhibition of Lipofuscin Accumulation in the Retina as a Therapeutic Strategy for Dry AMD Treatment. Drug Discov. Today Ther. Strateg. 2013, 10 (1), e11–e20. 10.1016/j.ddstr.2013.05.004.

(34) von Lintig, J. Metabolism of Carotenoids and Retinoids Related to Vision. J. Biol. Chem. 2012, 287 (3), 1627–1634. 10.1074/jbc.R111.303990.

(35) Cvekl, A.; Wang, W.-L. Retinoic Acid Signaling in Mammalian Eye Development. Exp. Eye Res. 2009, 89 (3), 280–291. 10.1016/j.exer.2009.04.012.

(36) Justilien, V.; Pang, J.-J.; Renganathan, K.; Zhan, X.; Crabb, J. W.; Kim, S. R.; Sparrow, J. R.; Hauswirth, W. W.; Lewin, A. S. SOD2 Knockdown Mouse Model of Early AMD. Invest. Ophthalmol. Vis. Sci. 2007, 48 (10), 4407–4420. 10.1167/iovs.07-0432.

(37) Jang, Y. P.; Matsuda, H.; Itagaki, Y.; Nakanishi, K.; Sparrow, J. R. Characterization of Peroxy-A2E and Furan-A2E Photooxidation Products and Detection in Human and Mouse Retinal Pigment Epithelial Cell Lipofuscin*. J. Biol. Chem. 2005, 280 (48), 39732–39739. 10.1074/jbc.M504933200.

(38) Radu, R. A.; Mata, N. L.; Nusinowitz, S.; Liu, X.; Sieving, P. A.; Travis, G. H. Treatment with Isotretinoin Inhibits Lipofuscin Accumulation in a Mouse Model of Recessive Stargardt’s Macular Degeneration. Proc. Natl. Acad. Sci. U. S. A. 2003, 100 (8), 4742– 4747. 10.1073/pnas.0737855100.

(39) Weng, J.; Mata, N. L.; Azarian, S. M.; Tzekov, R. T.; Birch, D. G.; Travis, G. H. Insights into the Function of Rim Protein in Photoreceptors and Etiology of Stargardt’s Disease from the Phenotype in Abcr Knockout Mice. Cell 1999, 98 (1), 13–23. 10.1016/S0092-8674(00)80602-9.

(40) Karan, G.; Lillo, C.; Yang, Z.; Cameron, D. J.; Locke, K. G.; Zhao, Y.; Thirumalaichary, S.; Li, C.; Birch, D. G.; Vollmer-Snarr, H. R.; Williams, D. S.; Zhang, K. Lipofuscin Accumulation, Abnormal Electrophysiology, and Photoreceptor Degeneration in Mutant ELOVL4 Transgenic Mice: A Model for Macular Degeneration. Proc. Natl. Acad. Sci. U. S. A. 2005, 102 (11), 4164–4169. 10.1073/pnas.0407698102.

(41) Maugeri, A.; Meire, F.; Hoyng, C. B.; Vink, C.; Van Regemorter, N.; Karan, G.; Yang, Z.; Cremers, F. P. M.; Zhang, K. A Novel Mutation in the ELOVL4 Gene Causes Autosomal Dominant Stargardt-like Macular Dystrophy. Invest. Ophthalmol. Vis. Sci. 2004, 45 (12), 4263–4267. 10.1167/iovs.04-0078.

