## SUPPLEMENTARY MATERIAL for "Rescue of the Stargardt Disease phenotype in *Abca4* knockout mice through dietary modulation of the vitamin A receptor RBPR2"

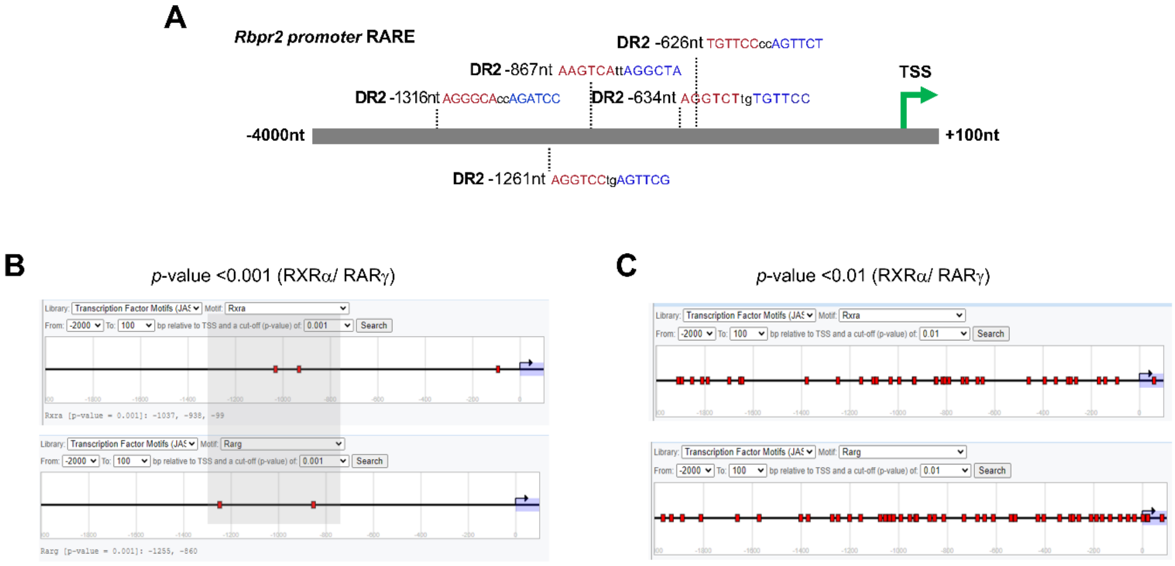


**Supplementary Figure S1: Computer based predictions of RARE sites on the murine *Rbpr2* gene promoter.**

Two publicly available resources, a software program and a server-based database; NUBIScan version 2.0 and EPD (Eukaryotic Promoter Database) were used (**A**) to identify putative nuclear receptor RARγ and RXRα (DR2 repeats) binding sites on the murine *Rbpr2* gene promoter. *Rbpr2* gene promoter analysis for the presence of RARγ and RXRα binding elements was further confirmed using the EPD-Eukaryotic Promoter Database and by setting statistically significant cut-off values at (**B**) $p<0.001$ and (**C**) $p<0.01$. DR2, direct repeat 2; RARE, retinoic acid response elements; TSS, transcription start site.

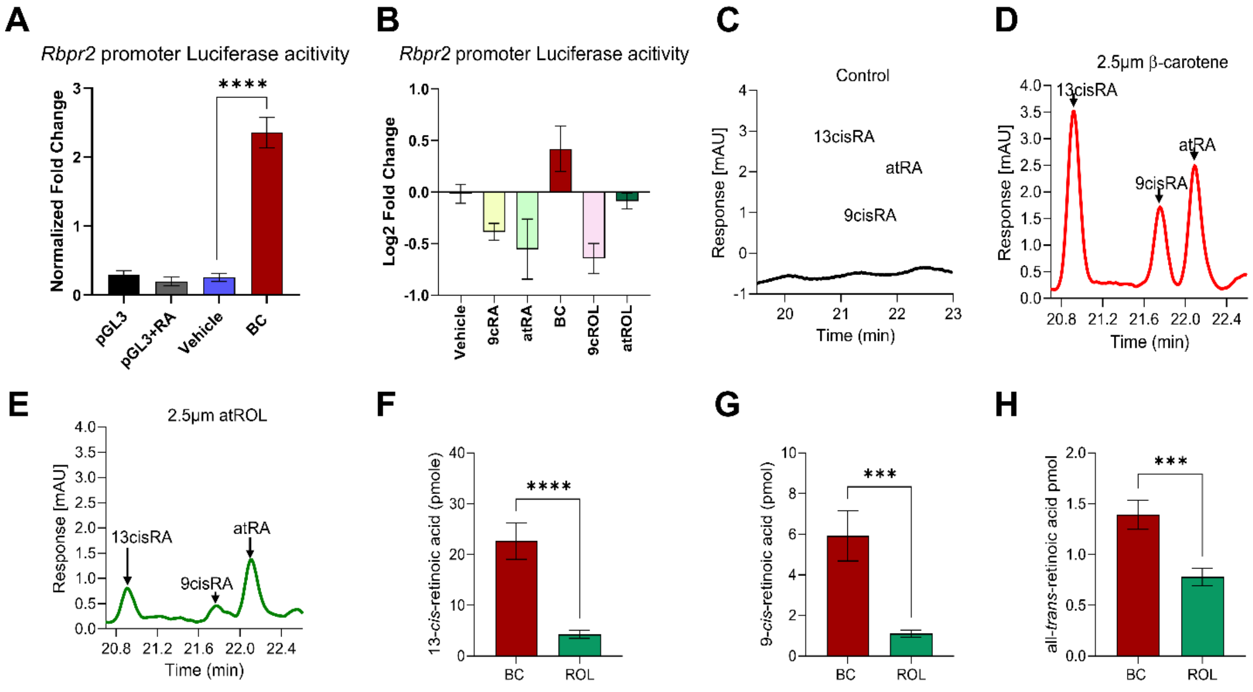


**Supplementary Figure S2: Exogenous β-carotene induces *Rbpr2* mRNA expression in vitro.**

**(A, B)** A firefly luciferase reporter assay was used to evaluate translational activity of the *Rbpr2* mRNA 5′-UTR region. COS1 cells were co-transfected with the wild-type (WT) *Rbpr2* 5′-UTR firefly reporter (pGL3-*Rbpr2*) and a control *Renilla* reporter and then treated with either all-*trans*-retinoic acid (atRA), 9-*cis*-retinoic acid (9cRA), β-carotene (BC), 9-*cis*-retinol (9cROL), all-*trans*-retinol (atROL), or vehicle (DMSO), all at 2 μM concentration for 24 h. *Rbpr2* mRNA promoter activity is represented by normalized fold change and Log2 fold change. (**C-E**) HPLC analysis of various retinoic acid isomers extracted from COS1 cells, post pGL3-*Rbpr2* promoter plasmid transfection and treatment with either BC or atROL. Control, are parental COS1 cells with no-treatment or transfection. (**F-H**) Quantification of various retinoic acid isomers in BC or atROL treated COS1 cells, post pGL3-*Rbpr2* promoter plasmid transfection. atRA, all-*trans*-retinoic acid; 9cisRA, 9-*cis*-retinoic acid; 13cRA, 13-*cis*-retinoic acid. One way ANOVA or Student t-test; *P<0.05; ***P<0.005; ****P<0.001.

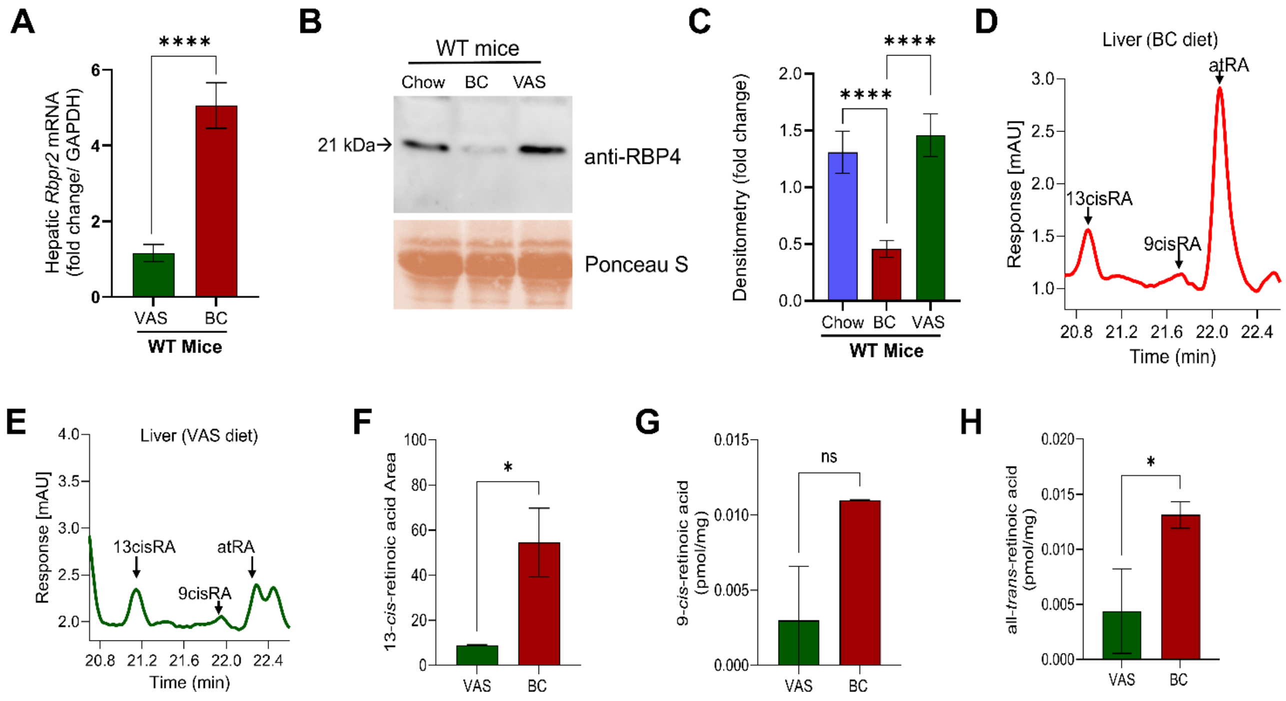


**Supplementary Figure S3: Dietary β-carotene supplementation induces hepatic *Rbpr2* mRNA expression and decreases serum RBP4 levels in WT mice.**

Cohorts of wild-type (WT) mice were fed a diet supplemented with β-carotene or a diet without β-carotene (VAS diet). Post 3-months of dietary intervention mice were euthanized and serum and liver harvested. (**A**) Quantitative RT-PCR analysis using *Rbpr2* gene specific probe set, was used to evaluate hepatic *Rbpr2* mRNA expression under the different dietary conditions. *Gapdh* mRNA expression was used as the internal control. (**B**) RBP4 protein expression in serum of mice determined by western blot analysis. Ponceau S was used to confirm equal protein loading among samples. (**C**) Densitometry quantification of serum RBP4 in cohorts of WT mice on different diets. (**D, E**) HPLC analysis of hepatic retinoic acid isomers extracted from mice fed a BC supplemented diet or a VAS diet devoid of BC. (**F-H**) Quantification of various retinoic acid isomers in liver of mice fed with dietary BC or a diet devoid of BC (VAS diet). atRA, all-*trans*-retinoic acid; 9cRA, 9-*cis*-retinoic acid; 13cRA, 13-*cis*-retinoic acid. One way ANOVA or Student t-test; *P<0.05; **P<0.01. n.s., not significant. n=3 mice per cohort.

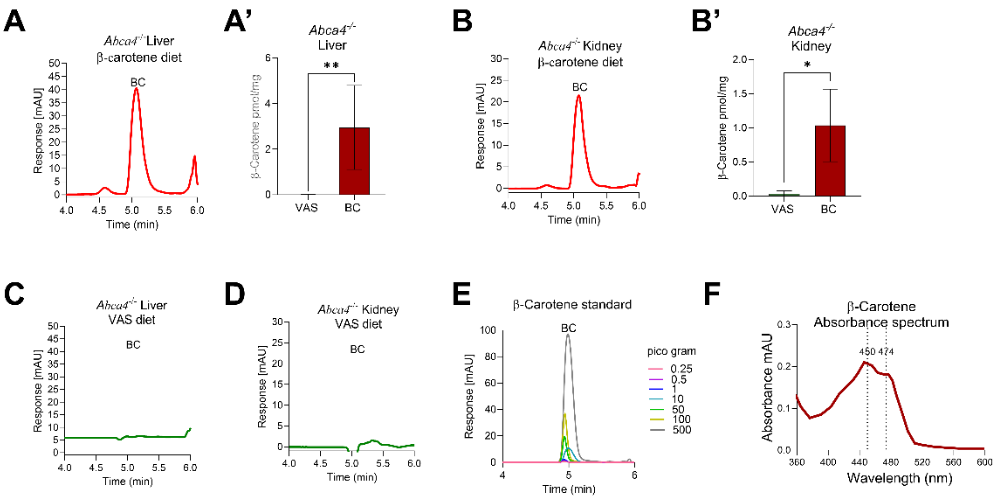


**Supplementary Figure S4: Quantification of β-carotene in mice tissue by HPLC.**

Approximately 100 mg of liver or kidney tissue from *Abca4*$^{-/-}$ mice fed either a β-carotene supplemented (**A, A', B, B'**) or VAS (**C, D**) diet was analyzed for β-carotene peaks using HPLC. (**E**) HPLC traces of β-carotene commercial standard at various concentrations and (**F**) β-carotene UV absorbance spectrum. BC, β-carotene; VAS, vitamin A sufficient diet. Student t-test; *P<0.05; **P<0.01.

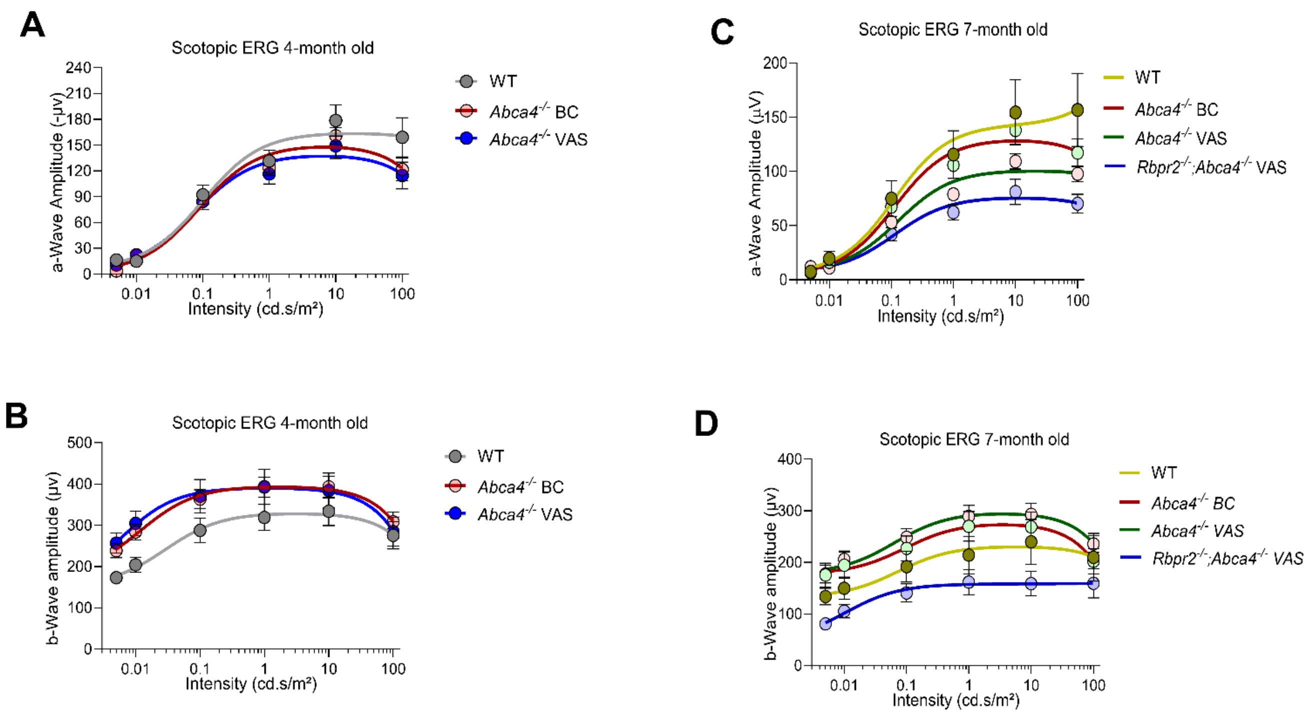


**Supplementary Figure S5: Scotopic ERG analysis in mice fed either a VAS or β-carotene supplemented diet.**

Dark-adapted scotopic electroretinographic (ERG) analysis in mice at 4- and 7-months of age, showing intensity range from 0.01 cd.s/m$^2$ to 100 cd.s/m$^2$. (**A**) *a*-wave amplitudes and (**B**) *b*-wave amplitudes at 4-months of age. (**C**) *a*-wave amplitudes and (**D**) *b*-wave amplitudes at 7-months of age. n=6 mice per cohort. VAS, vitamin A sufficient diet; BC, β-carotene supplemented diet.

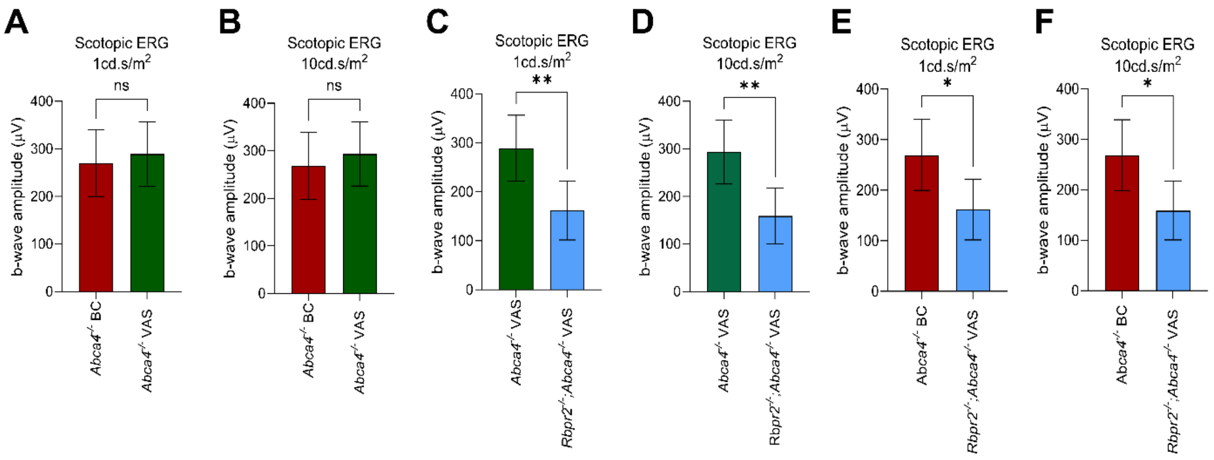


**Supplementary Figure S6: Scotopic *b*-wave analysis in mice fed either a VAS or β-carotene supplemented diet.**

Dark-adapted scotopic electroretinographic (ERG) *b*-wave analysis in various mice groups at 7-months of age, showing intensity range at 1 cd.s/m$^2$ (panels **A, C, E**) and 10 cd.s/m$^2$ (panels **B, D, F**). n=6 mice per cohort. VAS, vitamin A sufficient diet; BC, β-carotene supplemented diet. Student t-test; *P<0.05; **P<0.01. n.s., not significant.

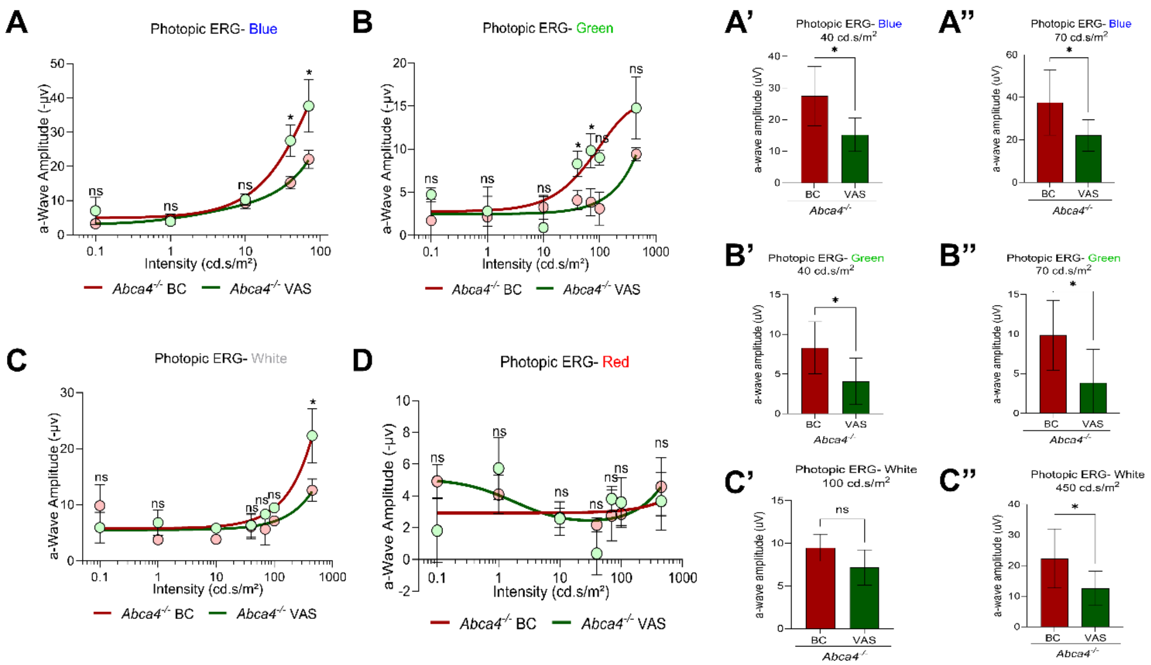


**Supplementary Figure S7: Long-term dietary β-carotene supplementation of *Abca4*$^{-/-}$ mice improves cone photoreceptor cell function.**

Light-adapted photopic electroretinographic (ERG) analysis in mice at 7-months of age. (**A, A', A''**) blue-light, (**B, B', B''**), green light, (**C, C', C''**) white light, and (**D**) red light, at various light intensities (40-, 70-, 100-, and 450- cd.s/m$^2$). n=4-6 mice per cohort. VAS, vitamin A sufficient diet; BC, β-carotene supplemented diet. n=6 mice per cohort. Student t-test; *P<0.05; n.s., not significant.

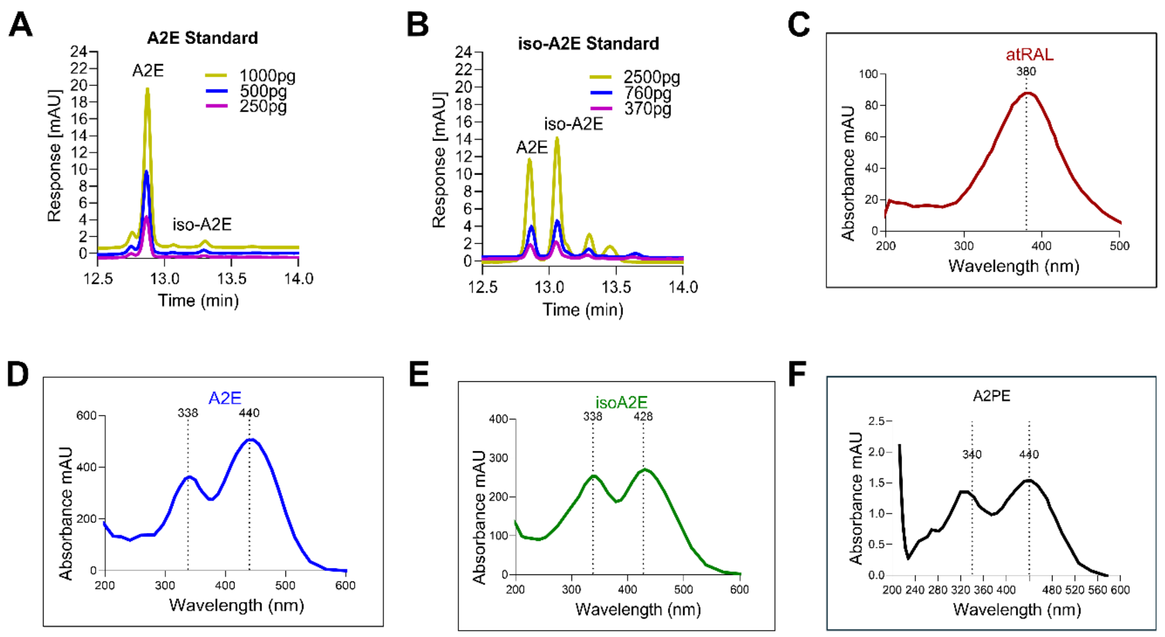


**Supplementary Figure S8: HPLC chromatograms and absorbance spectrum of ocular A2E isoforms.**

(**A, B**) HPLC traces of A2E and iso-A2E commercial standards at various concentrations. UV absorbance spectrum of (**C**) all-*trans*-retinal, (**D**) A2E, (**E**) iso-A2E, and (**F**) A2PE, from *Abca4*$^{-/-}$ mice eyes.
